# Discovery and structural analysis of glycoside hydrolase family 176 α-1,2 glucosidase from *Arthrobacter humicola* A8F5

**DOI:** 10.64898/2026.07.01.735942

**Authors:** Ryota Yasukochi, Takashi Suzuki, Terumasa Toraya, Keiko Hino, Tetsuya Mori, Toma Kashima, Akimasa Miyanaga, Hikaru Watanabe, Shinya Fushinobu

## Abstract

Glycoside hydrolases (GHs) exhibit remarkable specificity dictated by the structural configuration of their target glycosidic linkages. While enzymes that process α-1,4- and α-1,6-linkages in starch or glycogen are well-characterized, those acting on less common bonds, such as α-1,2-glucosidic linkages, remain largely underexplored. In this study, we report the discovery and structural elucidation of a novel α-1,2-glucosidase from *Arthrobacter humicola* A8F5 (A8F5 glucosidase), representing a newly uncovered activity within the poorly characterized GH176 family. Biochemical characterizations revealed that A8F5 glucosidase exclusively cleaves α-1,2-linkages via an anomer-inverting mechanism, with a distinct preference for short kojioligosaccharides. To circumvent crystallization obstacles caused by high loop flexibility and translational non-crystallographic symmetry, we engineered a loop-truncated variant. This strategy enabled the determination of high-resolution (up to 1.79 Å) crystal structures of the enzyme in its ligand-free form and in complex with kojibiose, kojitriose, and selaginose. A8F5 glucosidase adopts a (α/α)_6_-barrel fold characteristic of clan GH-G. Complementing the crystal structures with AlphaFold3 prediction demonstrated that two prominent active-site loops (loops 3 and 4) adopt a closed conformation that constricts the catalytic pocket, rendering the architecture suitable for short oligosaccharide recognition while restricting access to larger polymers. Furthermore, sequence similarity network analysis highlights vast, uncharacterized functional diversity within the GH176 family. These findings revealed that the GH176 enzyme recognizes and hydrolyses α-1,2-glucosidic bonds through a structural framework distinct from that of the previously known clan GH-L GH65 kojibiose hydrolase, expanding the known functional landscape of this enzyme group toward rare α-glucans.

## Introduction

Carbohydrates exhibit vast diversity due to the variety of their monosaccharide units and linkages (1). Consequently, enzymes that cleave these glycosidic linkages also exhibit high structural and functional diversity (2, 3). Even when limited to glucose disaccharides, α- or β-glucosidic linkages can be formed via the hydroxy groups at C1, C2, C3, C4, or C6, yielding 11 regio- and stereoisomers, such as maltose (Glc-α-1,4-Glc), cellobiose (Glc-β-1,4-Glc), and trehalose (Glc-α-1,1-α-Glc) (4). Among α-glucans, polysaccharides such as starch (composed of α-1,4- and α-1,6-linkages), dextran (composed of α-1,6-linkages), pullulan (a polymer of α-1,4-linked maltotriose units connected by α-1,6-linkages), and nigeran (composed of alternating α-1,3- and α-1,4-linkages) have been widely utilized in the food and related industries and extensively studied (5, 6). However, an α-glucan polymer mainly composed of α-1,2 linkages has not yet been identified in nature. Among shorter oligosaccharides, the α-1,2-linked disaccharide kojibiose (Glc-α-1,2-Glc) and its corresponding oligosaccharides, such as kojitriose and kojitetraose, have been detected in *koji* extract, *sake*, beer, honey, and molasses, albeit only in trace amounts (7, 8). Due to its resistance to oral fermentation, capacity to stimulate beneficial gut microbiota (8), and activity as an α-glucosidase inhibitor (9), kojibiose has attracted significant attention. However, because of its low abundance in nature, efficient methods for its large-scale production have been sought. Alongside chemical synthesis (10), enzymatic approaches for the production of kojibiose and kojioligosaccharides have been developed, leveraging the reverse phosphorolysis activity of kojibiose phosphorylase (EC 2.4.1.230) (11) or the transglycosylation activity of sucrose phosphorylase (EC 2.4.1.7) (12).

Although numerous enzymes that cleave α-glucosidic linkages are known, until recently, only four classes of enzymes capable of cleaving α-1,2-glucosidic linkages had been characterized: kojibiose phosphorylase (13), processing α-glucosidase I (EC 3.2.1.106) (14), glucosylgalactosylhydroxylysine glucosidase (EC 3.2.1.107) (15), and dextran α-1,2-debranching enzyme (EC 3.2.1.115) (16, 17). Of these, kojibiose phosphorylase is a phosphorolytic enzyme rather than a hydrolase, and both glucosylgalactosylhydroxylysine glucosidase and dextran α-1,2-debranching enzyme are specific to branched substrates, exhibiting negligible activity toward linear kojibiose and kojitriose. More recently, a dedicated kojibiose hydrolase (EC 3.2.1.216) that specifically hydrolyzes the α-1,2-glucosidic linkage of kojibiose was discovered in 2021 (18, 19). Among these five enzymes, processing α-glucosidase I belongs to glycoside hydrolase (GH) family 63 in the Carbohydrate-Active enZymes database (20), whereas the other four enzymes belong to GH65. Consequently, the diversity of α-1,2-glucosidic linkage-specific enzymes across the protein sequence landscape remains unexplored.

In this study, through extensive screening of soil-derived microorganisms, we discovered an enzyme that specifically hydrolyzes α-1,2-glucosidic linkages. Sequence analysis revealed that this enzyme belongs to GH176, which was recently established based on a single starch-acting enzyme (putatively cleaving the α-1,6 branch) (21) and remains relatively understudied. We also determined the crystal structures of this enzyme in complex with its substrates, thereby elucidating the structural and functional features of this novel α-1,2-glucosidase.

## Results and Discussion

### Screening, cloning, and biochemical properties of a novel α-1,2 glucosidase

During the screening process, approximately 1,000 bacterial strains were isolated from soil and subsequently cultured to prepare crude enzymes. The crude enzymes were reacted with α-linked glucobioses and then the reaction products were then analyzed by thin-layer chromatography (TLC). The crude enzymes derived from bacterial strain A8F5 hydrolyzed only kojibiose (Fig. 1*A*). The DNA sequence of an approximately 1.5-kb fragment obtained by PCR amplification of the 16S rRNA gene from strain A8F5 was subjected to an EzTaxon database search (22). The top four closely related type strains with the highest sequence identity (all > 98.8%) belonged to the genus *Arthrobacter*, and the 16S rRNA gene sequence of strain A8F5 exhibited 100% identity to that of *Arthrobacter humicola* KV-653^T^ (23). Therefore, strain A8F5 was taxonomically identified as *Arthrobacter humicola*. The enzyme from the extract of *A. humicola* A8F5 was purified according to the method described in the Experimental procedures section. Edman degradation revealed that the N-terminal sequence was TVQPGLH. Subsequent whole-genome analysis of *A. humicola* A8F5 identified an open reading frame (ORF) encoding a deduced 652-amino-acid (aa) protein, and its sequence starting from the second residue perfectly matched the determined N-terminal sequence. Additionally, a potential Shine-Dalgarno sequence was identified upstream of the start codon, and a putative transcription terminator sequence was found downstream of the stop codon. The ORF showed 29.6% sequence identity (217/734 aa) with the founding member of the GH176 family, amylo-α-1,6-glucosidase from *Thermococcus gammatolerans* STB12 (TgAMY) (21), indicating that this enzyme (hereafter referred to as A8F5 glucosidase) belongs to the GH176 family.

**Figure 1.**
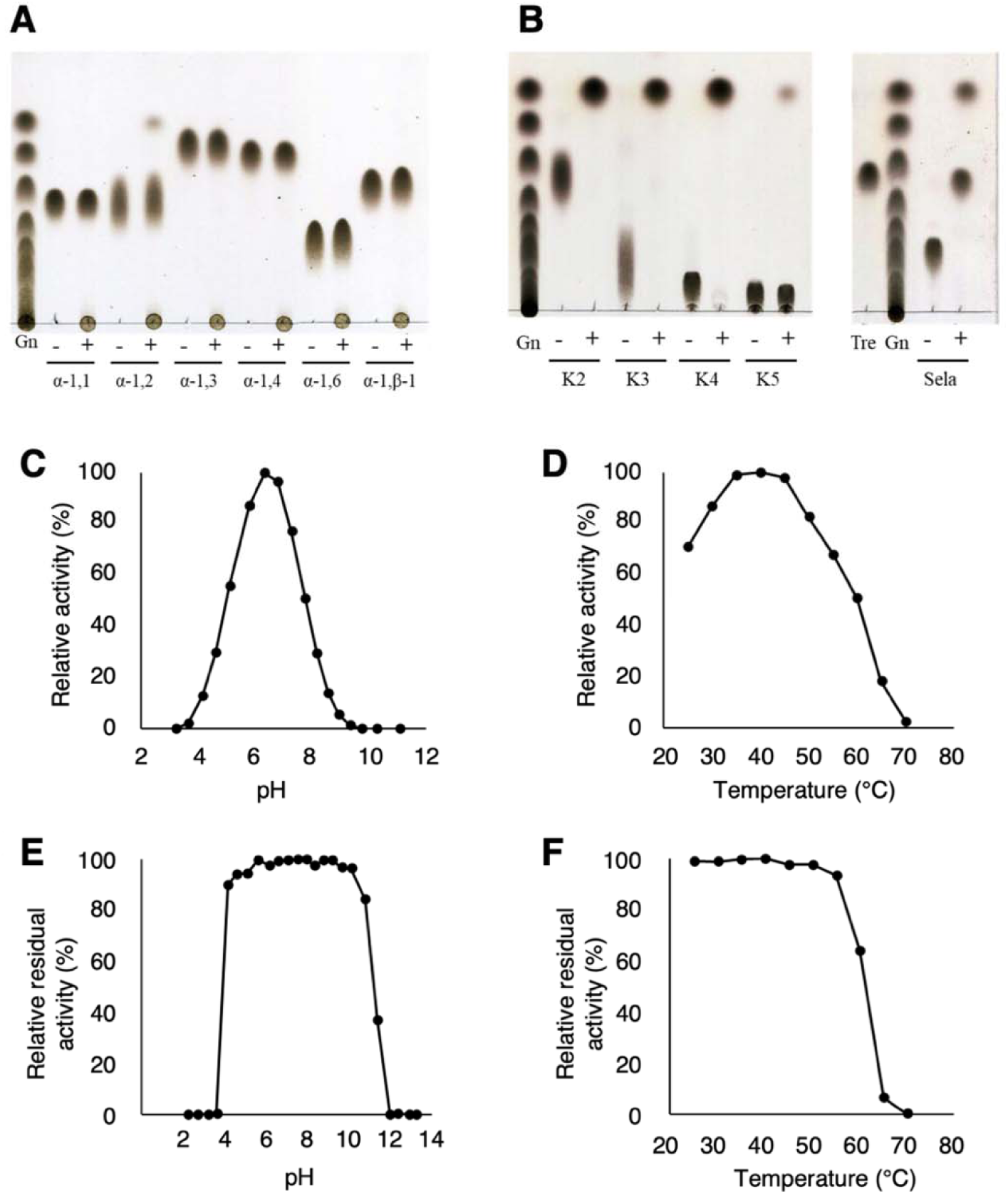
Enzymatic characteristics of A8F5 glucosidase. *A* and *B*, TLC analysis of the enzymatic activity. *A*, Hydrolytic activity of the crude enzyme from strain A8F5 toward various α-linked glucobioses. *B*, Hydrolytic activity of A8F5 glucosidase toward various α-1,2-linked oligosaccharides. The symbols are defined as follows: Gn, glucose and maltooligosaccharides; – and +, without or with the enzyme; α-1,1, trehalose; α-1,2, kojibiose; α-1,3, nigerose; α-1,4, maltose; α-1,6, isomaltose; α,-1,β-1, neotrehalose; K2, kojibiose; K3, kojitriose; K4, kojitetraose; K5, kojipentaose; Tre, trehalose; Sela, selaginose (2-O-α-glucosyl trehalose). *C*-*F*, Effects of pH and temperature on the activity and stability. *C*, Effect of pH on the activity measured in 50 mM Britton-Robinson buffer at 40°C. *D*, Effect of temperature on the activity at pH 6.5. *E*, pH stability measured by the residual activity after incubation in 50 mM Britton-Robinson buffer at various pH values and 4°C for 24 h. *F*, Thermal stability measured by the residual activity after incubation at various temperatures for 1 h. The standard assay condition was pH 6.5 at 40°C.

A8F5 glucosidase was recombinantly expressed as a His-tagged protein in *Escherichia coli*, and its activity was confirmed by TLC. The recombinant enzyme exhibited the same catalytic activity as the native enzyme. Regarding its enzymatic properties, including the effects of pH, temperature, and various metal ions on activity, A8F5 glucosidase showed maximal activity at pH 6.3 and 40°C (Fig. 1*C, D*) and was stable over a pH range of 4 to 10 and up to 55°C (Fig. 1*E, F*). The activity was not significantly affected by the addition of most metal ions, except for Cu² and Hg², which markedly inhibited the reaction (Table S1). Next, the substrate specificity for α-linked glucobioses was analyzed by high-performance liquid chromatography (HPLC). The enzyme strongly hydrolyzed kojibiose (specific activity 59.1 ± 0.1 U/mg), and weakly hydrolyzed nigerose (α-1,3 linkage; 0.05 ± 0.02 U/mg). In contrast, hydrolysis of trehalose (α,α-1,1 linkage), maltose (α-1,4 linkage), isomaltose (α-1,6 linkage) and neotrehalose (α,β-1,1 linkage) was not detected, demonstrating that A8F5 glucosidase is a highly specific α-1,2-glucosidase. The kinetic parameters for kojibiose and nigerose were measured (Table 1). Compared with those for nigerose, the *K*_m_, *k*_cat_, and *k*_cat_/*K*_m_ values for kojibiose were 86-fold lower, 306-fold higher, and 26,500-fold higher, respectively. The substrate specificity toward oligosaccharides with α-1,2 linkages was analyzed by TLC. A8F5 glucosidase effectively hydrolyzed kojibiose, kojitriose, and kojitetraose, while its activity toward kojipentaose was relatively low (Fig. 1*B*). Moreover, selaginose (2-*O*-α-glucosyl trehalose) was effectively degraded into trehalose and glucose. The anomeric configuration of the hydrolysis product released by A8F5 glucosidase was analyzed by gas-liquid chromatography. The chromatogram of the trimethylsilyl derivatives of the products indicated that the newly generated glucose formed from selaginose and kojibiose by the enzyme was the β-anomer, demonstrating that A8F5 glucosidase hydrolyzes the substrates via an anomer-inverting mechanism (Fig. S1).

**Table 1.** Kinetic parameters of A8F5 glucosidase for kojibiose and nigerose.

| Substrate | $K_m$ (mM) | $k_{cat}$ (s<sup>-1</sup>) | $k_{cat}/K_m$ (mM s<sup>-1</sup>) |
| --- | --- | --- | --- |
| Kojibiose | 1.15 ± 0.03 | 91.1 ± 1.7 | 79.5 ± 0.7 |
| Nigerose | 98.9 ± 3.9 | 0.30 ± 0.01 | 0.003 ± 0.00008 |

Kinetic parameters for kojibiose and nigerose were measured under the optimal condition (pH 6.5, 40°C). The amounts of glucose released by hydrolysis were determined by HPLC.

### Overall structure

The crystal structure of wild-type (WT) A8F5 glucosidase was determined at 1.90 Å resolution in the space group *P*2_1_2_1_2_1_ (Table S2), with a glucose molecule bound in the active site (Fig. 2*A*). A8F5 glucosidase forms a homodimer in the crystal, with the two monomers related by a non-crystallographic 2-fold symmetry. This dimeric state is consistent with results from size-exclusion chromatography (Fig. S2), indicating that the protein functions as a dimer in solution. The monomer of A8F5 glucosidase consists of two domains: an N-terminal β-sandwich domain and a C-terminal (α/α)_6_-barrel domain (Fig. 2*B*). The structure of A8F5 glucosidase features four prominent loop regions (loop 1 to loop 4) that exhibit distinct levels of flexibility in the dimer. In both chains, loop 2 (residues 239–243) is well-defined, whereas loop 1 (residues 33–58) and loop 3 (residues 339–350) are disordered, with no discernible electron density. Loop 4 (residues 412–421) exhibits asymmetry in the dimer, as it is disordered in chain A but clearly visible in chain B, likely due to differences in the local crystal packing environment. Loops 3 and 4 are proximal to the active site, whereas loops 1 and 2 are distal to it.

**Figure 2.**
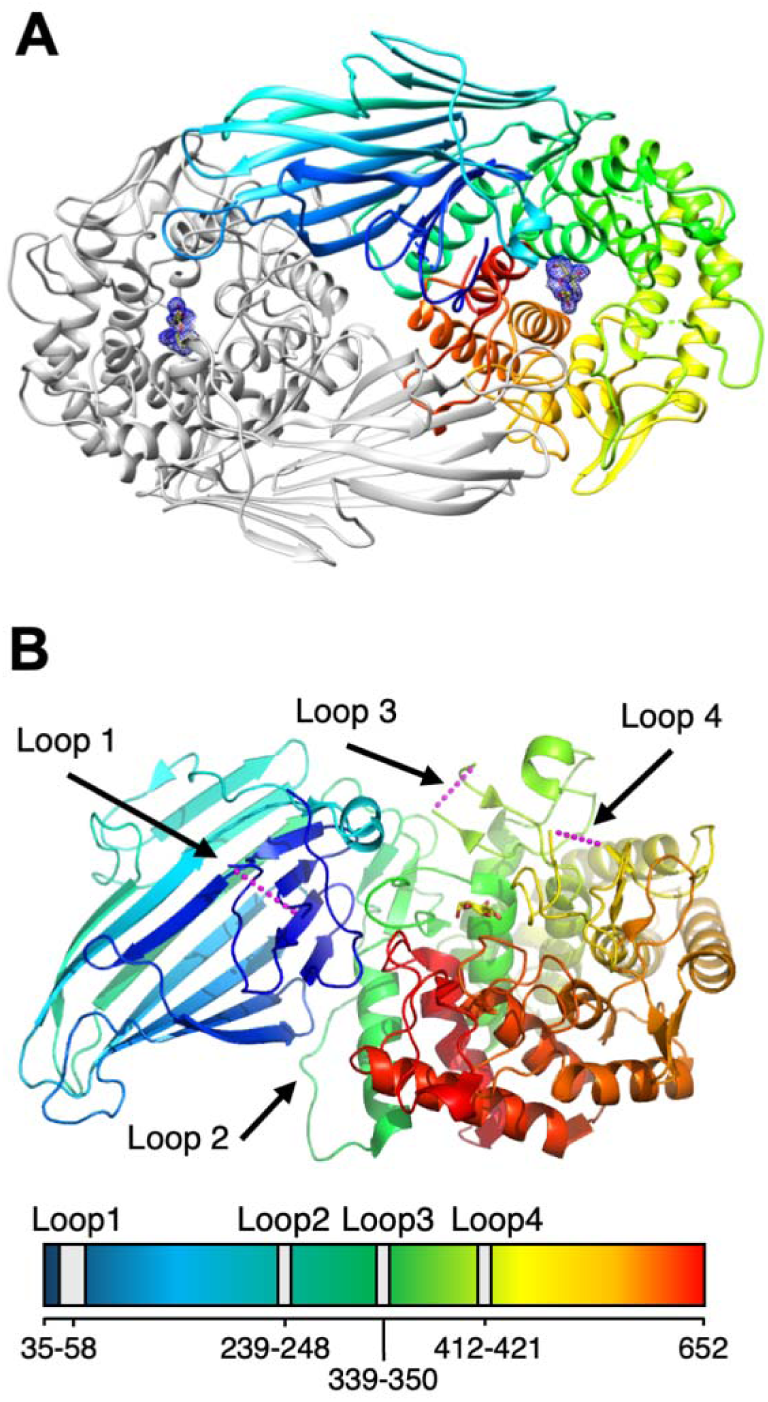
Overall structure and active site architecture of A8F5 glucosidase. *A*, Cartoon representation of the homodimeric structure of A8F5 glucosidase. One subunit is colored light gray, and the other is colored with a rainbow gradient from the N-terminus (blue) to the C-terminus (red). Polder maps for the bound glucose molecules are shown as blue meshes contoured at 3.0 σ. *B*, Monomeric structure of A8F5 glucosidase (chain A). The structure exhibits a two-domain architecture comprising an N-terminal β-sandwich domain and a C-terminal (α/α)_6_-barrel domain that houses the active site. The lower panel shows a schematic representation of the primary sequence and the loop distribution.

A Dali structural similarity search (24) revealed that A8F5 glucosidase exhibits moderate structural similarity to well-characterized, anomer-inverting α-glucosidase families: *e.g.*, GH100 “invertases” (sucrose-acting α-glucosidases rather than β-fructofuranosidases) (25, 26), GH63 YgjK (27), and GH37 trehalases (28) (Table 2). Because the GH37, GH63, and GH100 families adopt an (α/α)_6_-barrel fold and constitute clan GH-G (20), they share a conserved set of catalytic residues, specifically the general acid and base. Although the sequence identity between A8F5 glucosidase and the GH-G enzymes was very low (< 18%), structural comparison enabled us to infer the catalytic residues (Fig. 3*A–D*). D427 and E611 in A8F5 glucosidase were suggested to be the catalytic acid and base residues, respectively, as they correspond to D501 and E727 of GH63 YgjK, for example. The carboxyl groups of D427 and E611 are separated by a distance of 8.6 Å, well within the typical range for α-anomer-inverting GHs (6.0–10.2 Å) (29).

**Figure 3.**
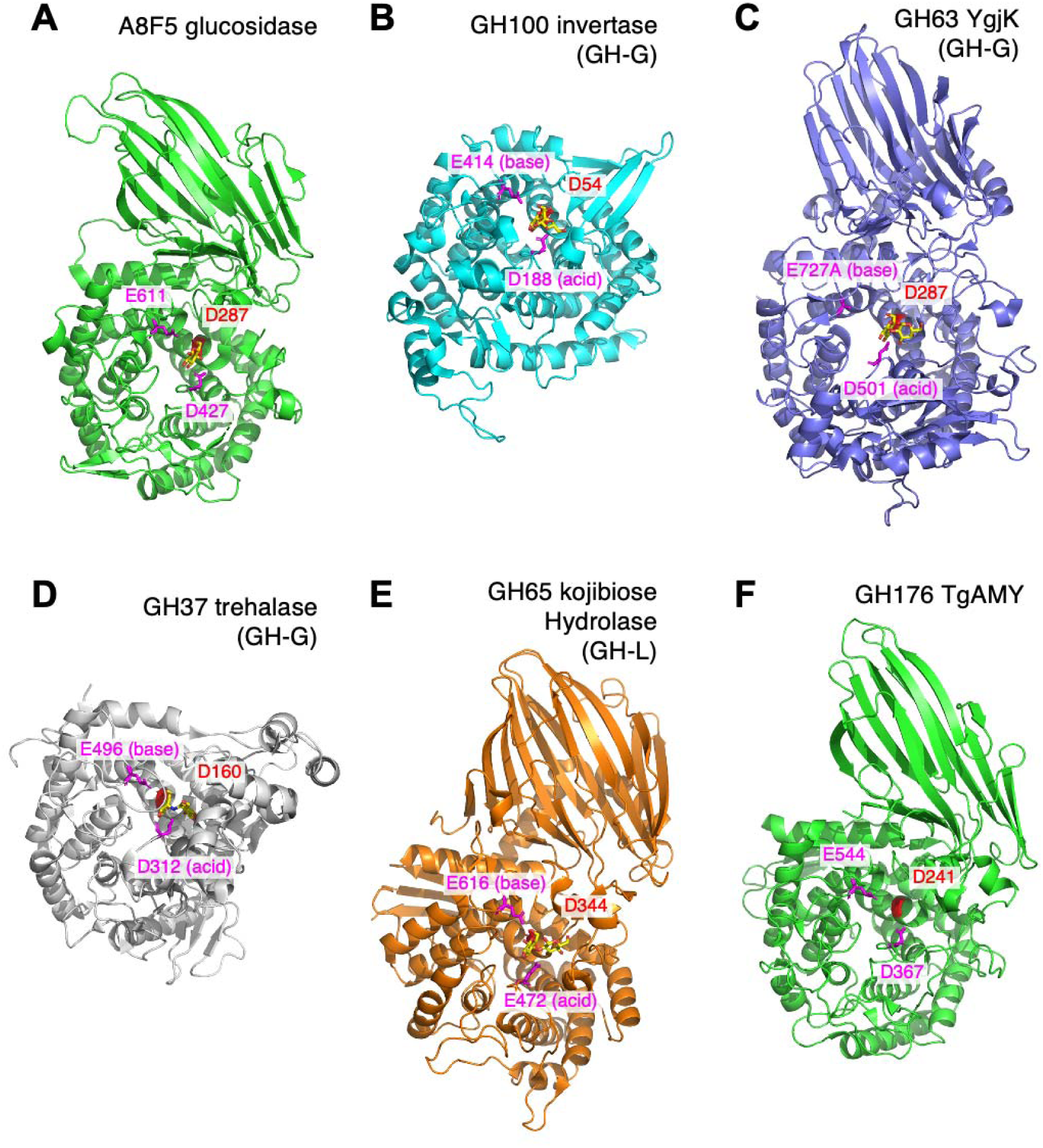
Comparison of the overall structure of A8F5 glucosidase with those of other related α-glucosidases. *A*, Wild-type A8F5 glucosidase complexed with glucose. *B*, GH100 invertase complexed with sucrose (PDB ID: 5GOP). *C*, GH63 α-glucosidase E727A mutant complexed with Glc-α-1,2-Gal (PDB ID: 3W7W). *D*, GH37 trehalase complexed with validoxylamine A (PDB ID: 2JF4). *E*, GH65 kojibiose hydrolase FjGH65A complexed with glucose (PDB ID: 7FE4). *F*, GH176 TgAMY (PDB ID: 7VMA). The conserved catalytic acid (D427, magenta), catalytic base (E611, magenta), and fixer (D287, red) residues in A8F5 glucosidase, as well as the corresponding residues in the other enzymes are shown as sticks.

**Table 2.**
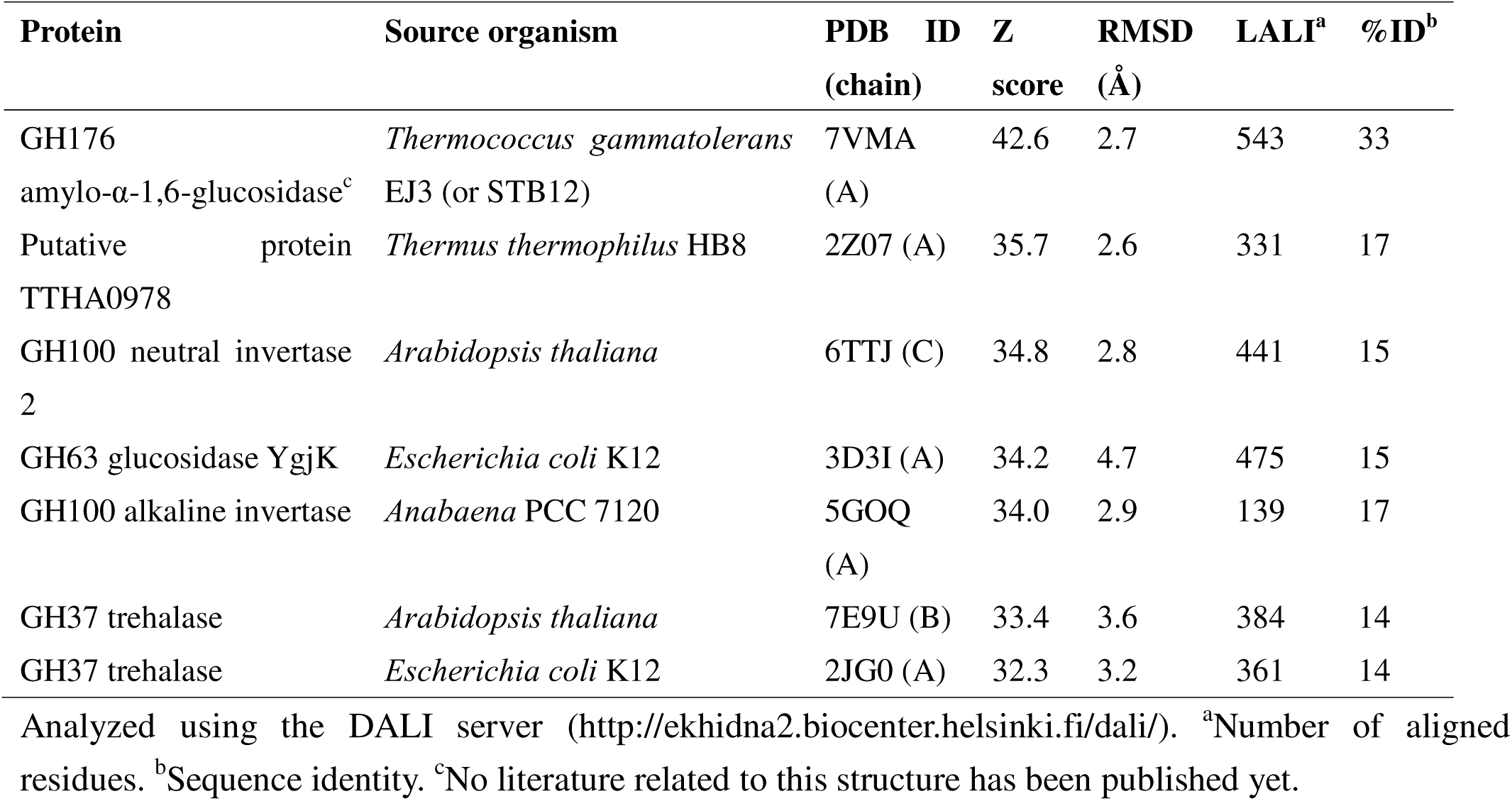
Results of DALI structural similarity search.

### Loop deletion construct

Initially, we attempted to determine the complex structures with uncleaved substrates using catalytically inactive D427N or D427A mutants. However, these attempts were unsuccessful, as the resulting crystals yielded only poor X-ray diffraction that was insufficient for reliable molecular replacement and subsequent model building. Analysis using phenix.xtriage revealed that the WT crystals exhibited strong translational non-crystallographic symmetry (tNCS) (30), with an off-origin Patterson peak at fractional coordinates (0.030, 0.000, 0.500) and a height of approximately 47% relative to the origin (p = 9.9 × 10□□). This suggests that the lattice of the WT crystal suffered from packing degeneracy, likely due to the high intrinsic flexibility of surface-exposed loops, which allowed slight translational offsets between neighboring dimers and hindered the formation of a uniquely ordered crystal lattice. To overcome this challenge, we designed 15 loop-deleted candidates by replacing the longest flexible loop (loop 1) with various short linker sequences (Table S3). These candidates were then screened *in silico* using AlphaFold3 (31) to evaluate their structural stability and local folding (Fig. 4*A*). The deletion variant exhibiting the highest predicted local distance test (pLDDT) score among these candidates, where the 26 amino acids from N33 to A58 were replaced with a short linker (SGS), was selected as Δloop1. The Δloop1 protein construct showed a protein expression level and catalytic activity similar to those of the WT enzyme (Fig. S2), and the purified protein crystallized with high reproducibility.

**Figure 4.**
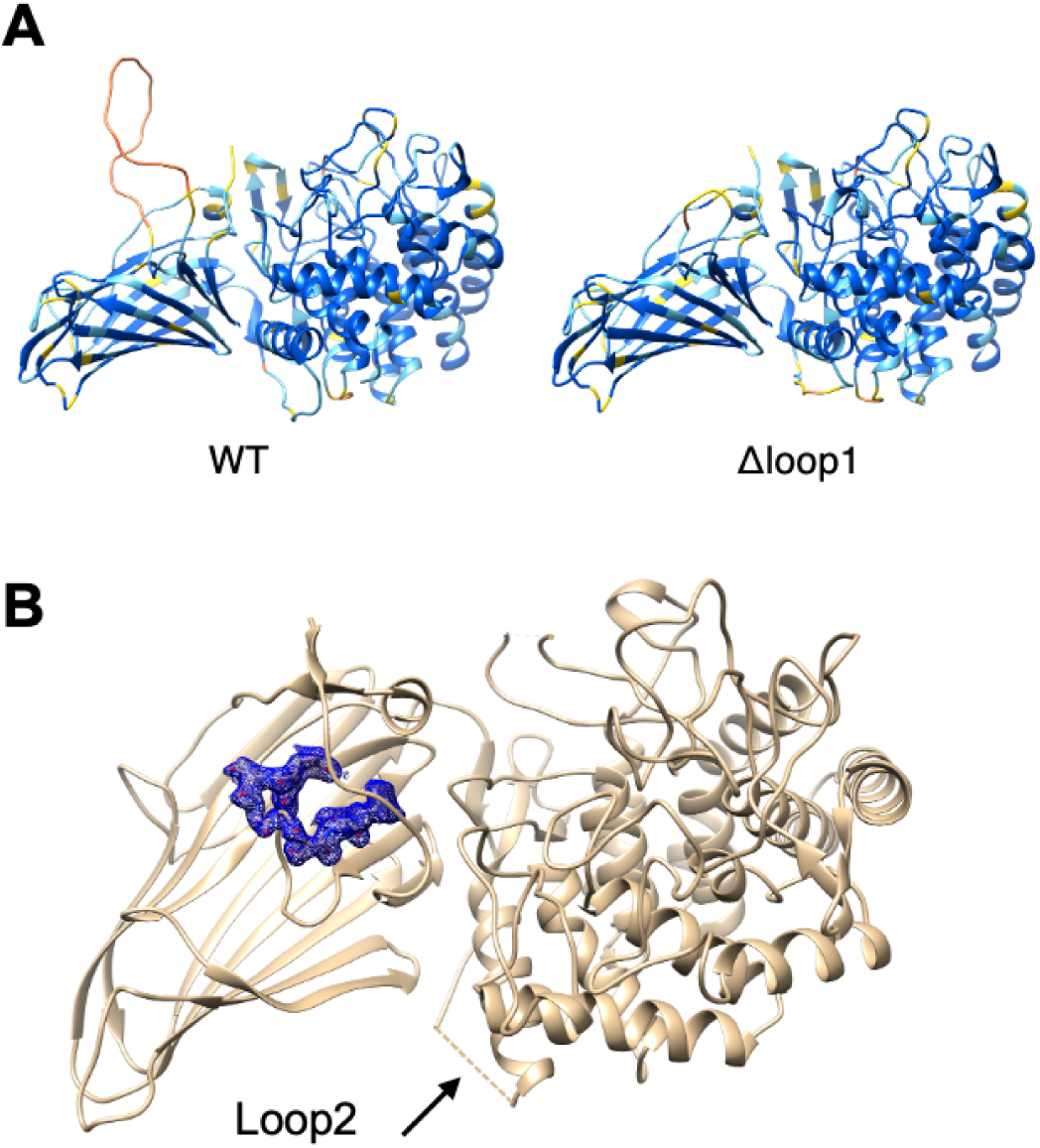
Structure-based design and validation of the Δloop1 construct. *A*, AlphaFold3-predicted structures of the WT (left) and Δloop1 (right) colored by pLDDT from low (orange) to high (blue). *B*, The ligand-free crystal structure of the Δloop1 construct. Polder map for the engineered linker region (GESGGSGF) is shown as a blue mesh contoured at 3σ. In the Loop 2 region, the electron density was too weak to build a model.

The ligand-free structure of Δloop1 was determined at 1.88 Å resolution in the space group *P*2_1_, and the asymmetric unit contained a dimer (Table S2). Notably, no significant tNCS was detected in the Δloop1 crystal (maximum Patterson peak height of ∼4.8%, p = 0.95), indicating that shortening the long flexible loop effectively reduced packing degeneracy and favored a more rigidly ordered crystal lattice. The overall structure of Δloop1 remained essentially identical to that of the WT enzyme, with a Cα root mean square deviations (RMSD) of 0.66 Å, confirming that the loop truncation did not perturb the global fold or the catalytic pocket architecture. The Polder map for the shortened loop 1, including the three-amino-acid linker, clearly defined the conformation of this region (Fig. 4*C*). While the modification of loop 1 stabilized the crystal lattice, the electron density for loop 2 (residues R239-A243) became poorly defined, likely due to subtle changes in the local packing environment. However, because loop 2 is located distal to the active site, this local flexibility was not expected to interfere with the substrate binding. These results validated the use of the Δloop1 construct as a reliable variant for subsequent structural studies of substrate complexes.

### Substrate recognition

The crystal structure of the Δloop1 D427N mutant in complex with kojibiose was determined at 1.79 □ resolution (Table S2). Clear electron density for kojibiose was observed in the catalytic pockets in the two chains (Fig. 5*A*–*B*). Kojibiose adopted slightly different binding conformations between the two chains. In chain A, the nonreducing-end glucoside at subsite –1 forms direct hydrogen bonds with the side chains of R286, D287, N427, H566, and Q634 and the main chain carbonyl of W425, while the reducing-end glucoside at subsite +1 (α-anomeric form) forms hydrogen bonds with the side chains of H335 and E336 (Fig. 5*A*). In particular, the substituted catalytic acid, N427, forms two hydrogen bonds with the O2 hydroxy group of the subsite –1 glucoside and the scissile glycosidic bond oxygen between subsites –1 and +1. D287 recognizes the C4 and C6 hydroxy groups of the glucoside at subsite −1 through two hydrogen bonds. This position corresponds to a residue that is highly conserved in GH15 glucoamylases (32) and in many glucoside-acting (α/α)_6_-barrel enzymes (26, 33, 34). Given that this glucoside-anchoring residue is conserved across GH families to a degree comparable to that of the two catalytic residues, we refer to it as the “fixer”, following the terminology used for GH13 α-amylases (35, 36). In addition to these hydrogen bonds, a hydrophobic platform provided by the side chains of three tryptophan residues (W425, W571, and W636) stabilizes the sugar–protein interaction at subsite –1. In chain B, while the overall position and interactions at subsite –1 are similar to those in chain A, two water-mediated hydrogen bonds are formed between kojibiose and the side chains of Q634 and E616 (catalytic base, Fig. 5*B*). The reducing-end glucoside at subsite +1 is shifted relative to the position observed in chain A. Consequently, the reducing-end glucoside loses its hydrogen bonds with H335 and E336, forming a new hydrogen bond with N427 instead. To verify whether this alternative conformation was a consistent feature rather than a localized packing artifact, we analyzed multiple independent crystallographic datasets. Our analysis revealed that kojibiose can adopt both orientations, with both chain A- and chain B-type conformations observed across different crystals. Although the hydrogen bonds between the glucosyl moiety at subsite +1 and H335 or E336 are not sufficiently strong to be maintained consistently in all crystal forms, they likely contribute to the substrate preference of this enzyme for kojibiose over other α-glucobioses.

**Figure 5.**
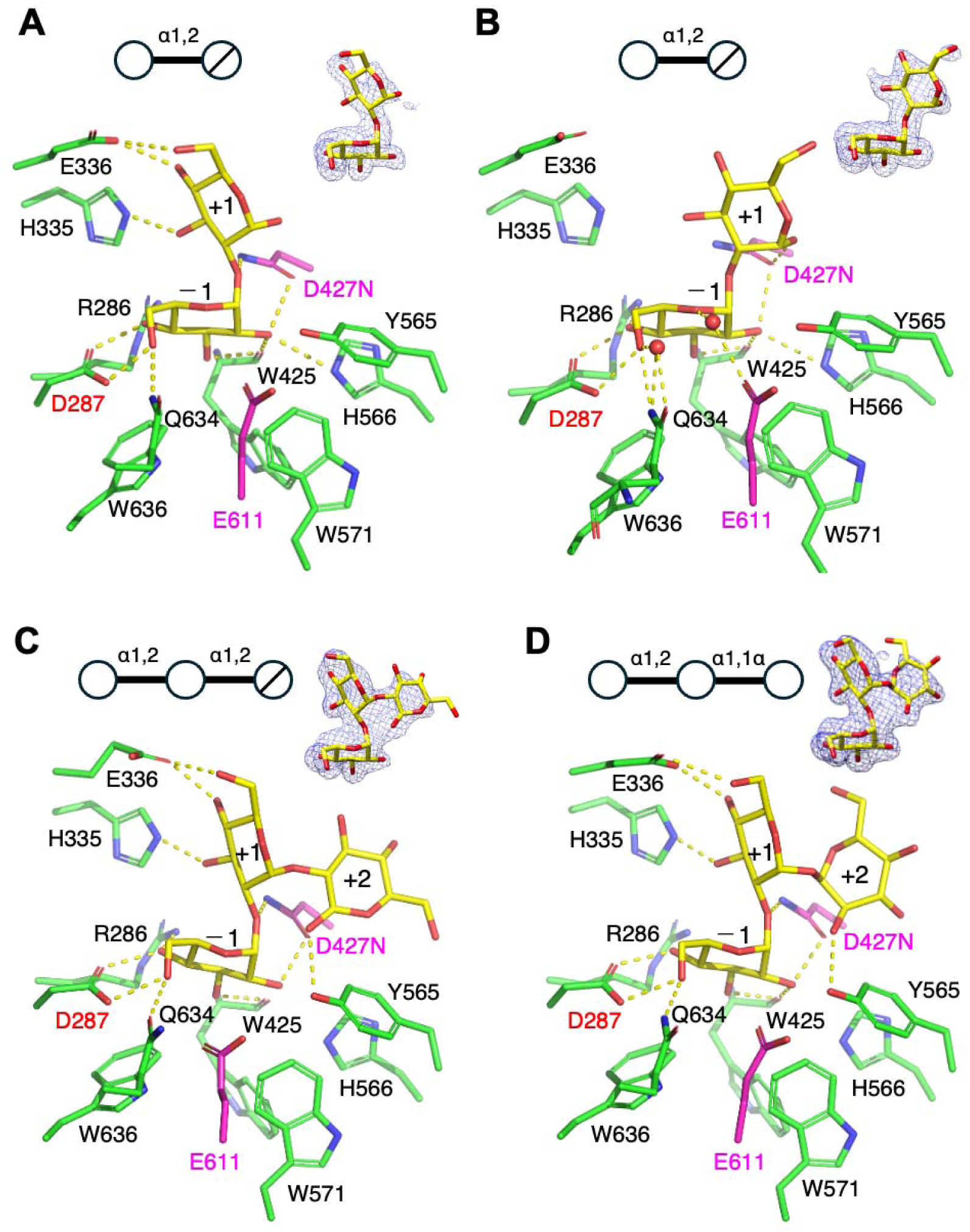
Active site structures of the inactive D427N mutant of A8F5 glucosidase Δloop1 construct in complex with the substrates. *A*, Chain A of the kojibiose complex. *B*, Chain B of the kojibiose complex. *C*, Chain B of the kojitriose complex. D, Chain B of the selaginose complex. Schematic representations of the ligand sugars, where a circle with a diagonal slash indicates a reducing-end sugar, and polder maps for the ligands (contoured at 4.0 σ for kojibiose and 3.5 σ for kojitriose and selaginose) are shown above each panel. Hydrogen bonds are shown as dashed lines.

Furthermore, we determined the complex structures in complex with kojitriose and selaginose at resolutions of 1.83 Å and 1.80 Å, respectively (Table S2). Figs. 5*C* and 5*D* display chain B because the binding mode in chain A was essentially identical, indicating that these trisaccharides are stably accommodated within the active sites of both subunits. Both kojitriose and selaginose consistently adopted the hydrogen-bond-rich orientation at subsite +1 that was observed for chain A in the kojibiose complex. The glucosides at subsites –1 and +1 of kojitriose and selaginose retained interaction patterns similar to those observed in the kojibiose complex. The glucoside at subsite +2 is linked via α-1,2- and α-1,1-α-linkages in kojitriose and selaginose, respectively, resulting in a flipped orientation of the sugar ring. The electron density revealed that the reducing-end glucoside of kojitriose adopts the β-anomeric configuration. The O1 hydroxy of this glucoside and the O2 hydroxy of selaginose are positioned similarly, and both form hydrogen bonds with Y565. The absence of stringent recognition at subsite +2 provides a plausible explanation for the ability of this enzyme to act on several kojibiose-containing oligosaccharides (Fig. 1*B*).

### Structural comparison with other α-glucosidases

As mentioned above, A8F5 glucosidase exhibits structural similarity to the α-glucosidases in GH100, GH63, and GH37, which belong to clan GH-G (Table 2). Because A8F5 glucosidase shares a common (α/α)_6_-barrel catalytic domain with these families and the positions of the two catalytic residues and the fixer residue are also conserved (Fig. 3*A*–*D*), GH176 may also be classified into clan GH-G. While GH176 and GH63 are two-domain enzymes consisting of an N-terminal β-sandwich domain and a C-terminal (α/α)_6_-barrel domain, GH100 and GH37 are single-domain enzymes lacking any associated domains. Enzymes in GH65, which include kojibiose hydrolase and kojibiose phosphorylase, also consists of two domains. However, the relative orientation of its β-sandwich and (α/α)_6_-barrel domains differs from that of GH176 and GH63 (Fig. 3*E*), and GH65 did not appear in the top 100 hits of the Dali structural similarity search. Accordingly, GH65 is grouped into clan GH-L along with GH15, GH125, and GH178.

The active site of A8F5 glucosidase was compared with those of other structurally similar GH family enzymes (Fig. 6). In clan GH-G enzymes such as GH100 invertase (26), GH63 YgjK (27), and GH37 trehalase (28), the positions of the two catalytic residues (aspartate as the acid and glutamate as the base) and the aspartate acting as the fixer, as well as their interactions with the glucoside at subsite -1, are almost fully conserved (Fig. 6*B–D*). Furthermore, two of the three tryptophan residues that form the hydrophobic platform in A8F5 glucosidase are also conserved (for example, W376 and W434 in GH100 invertase). However, other active-site residues are not conserved, and the interactions at subsite +1 are completely different. Because GH100, GH63, and GH37 enzymes preferentially hydrolyze sucrose, nigerose, and trehalose, respectively, this structural diversity in the active site, especially at subsite +1, reflects their distinct substrate specificities.

**Figure 6.**
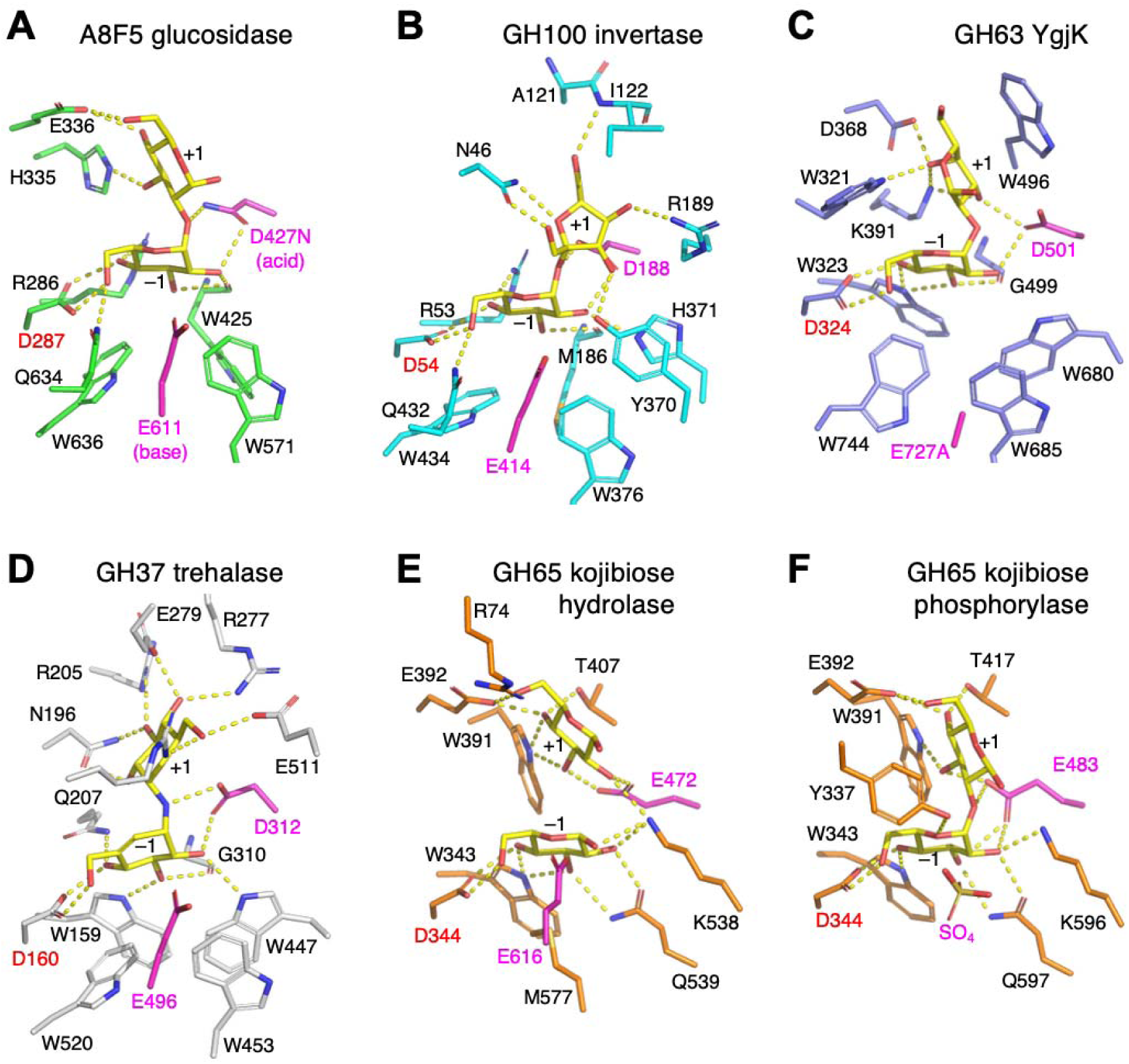
Comparison of the active site of A8F5 glucosidase with those of other related α-glucoside-cleaving enzymes. *A*, A8F5 glucosidase D427N mutant complexed with kojibiose. *B*, GH100 invertase complexed with sucrose (PDB ID: 5GOP). *C*, GH63 α-glucosidase E727A mutant complexed with Glc-α-1,2-Gal (PDB ID: 3W7W). *D*, GH37 trehalase complexed with validoxylamine A (PDB ID: 2JF4). *E*, GH65 kojibiose hydrolase FjGH65A complexed with glucose (PDB ID: 7FE4). *F*, GH65 kojibiose phosphorylase complexed with kojibiose and sulfate (PDB ID: 3WIQ). The catalytic residues are shown as magenta sticks.

In the two types of GH65 enzymes that utilize kojibiose as a substrate, namely kojibiose hydrolase FjGH65A from *Flavobacterium johnsoniae* (18) and kojibiose phosphorylase from *Caldicellulosiruptor saccharolyticus* (34), the general acid catalytic residue is glutamate instead of aspartate (Fig. 6*E*, *F*). Furthermore, kojibiose phosphorylase lacks a general base catalyst and instead possesses a phosphate-binding site, where a sulfate ion was bound in the crystal structure. These findings indicate that GH65 enzymes share only minimal commonalities with A8F5 glucosidase regarding the interactions involved in kojibiose recognition, as they utilize a fundamentally different structural framework. Interestingly, the ‘fixer’ aspartate residue is strictly conserved among all of the enzymes shown in Fig. 6.

### Structural comparison with another GH176 enzyme

The top hit in the Dali structural similarity search was the ligand-free crystal structure of GH176 TgAMY (PDB ID: 7VMA), but the paper regarding this structure has not yet been published. Superimposition of the active sites of the A8F5 glucosidase and TgAMY structures revealed that not only the catalytic residues but also all residues involved in kojibiose recognition are completely conserved (Fig. 7*A*). This finding is inconsistent with previous reports indicating that TgAMY acts on α-1,6-glucosidic bonds of maltodextrin (21).

**Figure 7.**
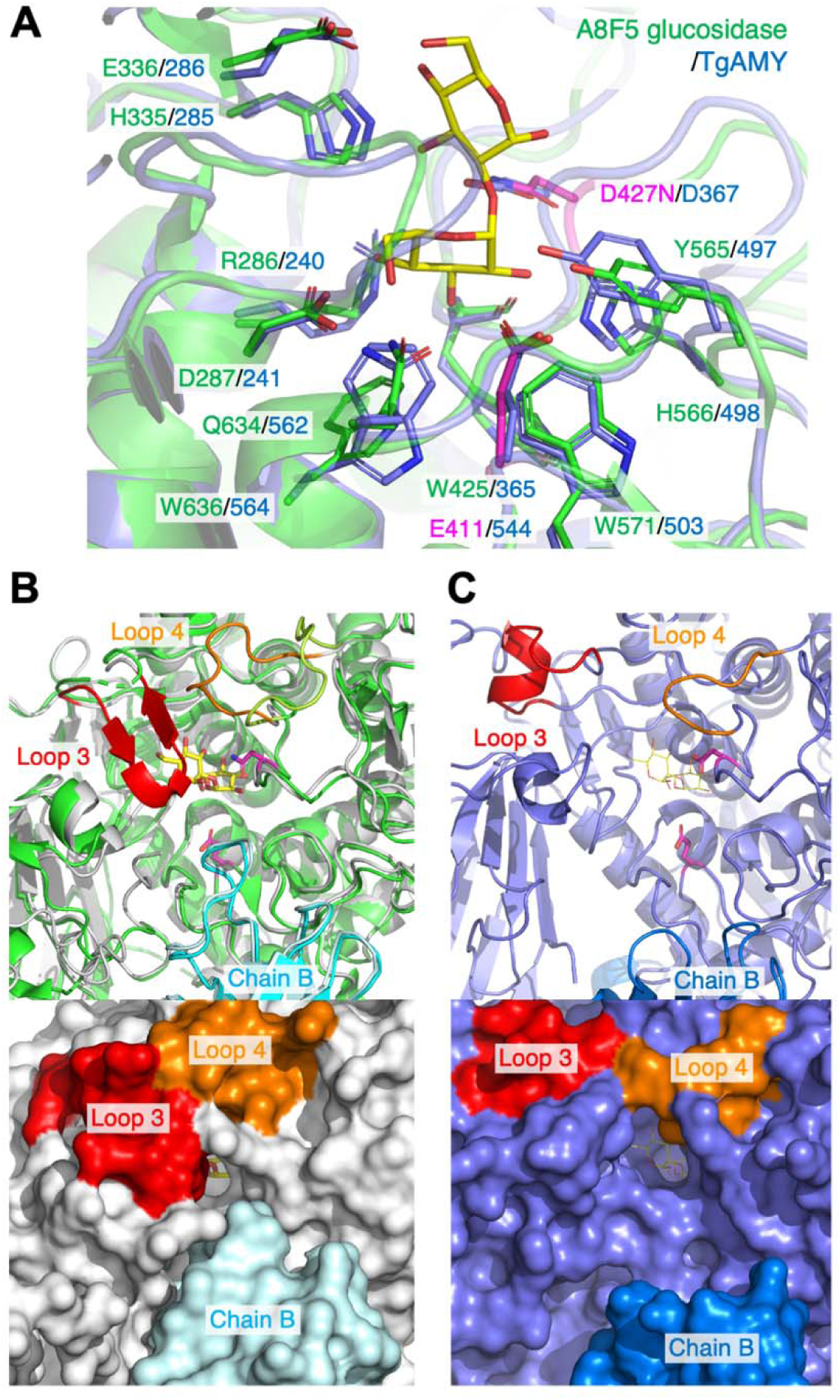
Structural comparison of A8F5 glucosidase with GH176 TgAMY. *A*, Superimposition of the active sites of A8F5 glucosidase (green) complexed with kojibiose (yellow) and TgAMY (blue). *B*, Active site structure of A8F5 glucosidase. Upper panel, the crystal structure of the kojibiose complex (green for chain A, cyan for chain B, and yellow for kojibiose) is superimposed with a structure predicted by AlphaFold3 (white for chain A, pale cyan for chain B, red for loop 3, and orange for loop 4). Lower panel, a surface model of the AlphaFold3-predicted structure showing the blocked substrate-binding pocket. *C*, Active site structure of TgAMY (blue for chain A, aquamarine for chain B, red for loop 3, and orange for loop 4). The position of kojibiose from A8F5 glucosidase is overlayed as thin yellow lines for reference. Lower panel, a surface model showing the open substrate binding site.

Because loop 3 was consistently disordered in the crystal structures of A8F5 glucosidase (Fig. 2*B*), structure prediction was performed using AlphaFold3 (31). The results predicted that while loop 3 does not directly interact with the substrate, it forms a short two-stranded β-sheet that caps over the active site (Fig. 7*B*, upper panel). In this structure, the adjacent loop 4 is predicted to adopt a closed conformation. Consequently, these two loops, together with the neighboring chain B, render the substrate-binding pocket of A8F5 glucosidase almost entirely blocked (Fig. 7*B*, lower panel). This narrow substrate-binding pocket environment aligns well with the experimental data showing that the activity of A8F5 glucosidase toward kojipentaose is significantly lower than its activity toward shorter substrates (Fig. 1*B*).

In contrast, both loop 3 and loop 4 of TgAMY are relatively short, with the former forming a short α-helix, and chain B is positioned away from the active site (Fig. 7*C*, upper panel). As a result, the substrate-binding site of TgAMY exhibits a more open structure (Fig. 7*C*, lower panel). This structural feature appears to match the reported characteristics of TgAMY, namely its preference on DE6 maltodextrin, which has a theoretical degree of polymerization of 17.7 (37), as well as larger polysaccharides like starch and glycogen (21). However, the fact that the active site of TgAMY forms a pocket with a blocked bottom (a feature typical of *exo*-type enzymes) contradicts the characterization of this enzyme as a debranching enzyme that cleaves branched α-1,6-glucosidic bonds (21). Debranching enzymes that preferentially cleave the α-1,6-branched chains of maltodextrins and starch, namely amylo-α-1,6-glucosidase (EC 3.2.1.33) and isoamylase (EC 3.2.1.68), predominantly belong to the GH13_11 subfamily and share common structural characteristics. Specifically, *Sulfolobus solfataricu*s TreX (38), *Escherichia coli* GlgX (39), and *Chlamydomonas* isoamylase 1 (40) all have a cleft-like substrate-binding site featuring an open plus-side subsite to accommodate branched chains. Notably, while indirect assays such as an increase in iodine staining, changes in chain length distribution, and the β-amylolysis limit analysis have been conducted on TgAMY, direct experimental evidence demonstrating the cleavage of α-1,6-glucosidic bonds was not provided (21). Therefore, further verification is required to determine exactly what specific catalytic activities exist within the GH176 family besides kojibiose hydrolase.

### Sequence similarity network (SSN) analysis

To investigate the distribution of proteins with amino acid sequences similar to that of A8F5 glucosidase, an SSN analysis was performed using 8,177 homologs available in the GenBank database (Fig. 8*A*). In the top 1,000 hits of the BLAST search with the lowest E-values, which are estimated to cover approximately 20% of the 4,892 members currently listed in GH176 (https://www.cazy.org/GH176.html), almost all (963 ORFs) are annotated as amylo-α-1,6-glucosidase or glycogen debranching enzyme, presumably following the report on TgAMY. At an alignment score cutoff of 100, both A8F5 glucosidase and TgAMY were partitioned into small, distinct clusters. Multiple sequence alignment of the loop 3 region, which is predicted to cap the active site in A8F5 glucosidase, revealed that it is highly conserved within the same cluster, whereas it is noticeably shorter in TgAMY (Fig. 8*B*). This indicates that each cluster shown in this SSN is classified in accordance with an enzyme function such as substrate specificity. Interestingly, at least six larger clusters of unknown function emerged in this SSN. Therefore, these results strongly suggest that GH176 potentially contains a large number of enzymes with functions distinct from those of A8F5 glucosidase and TgAMY.

**Figure 8.**
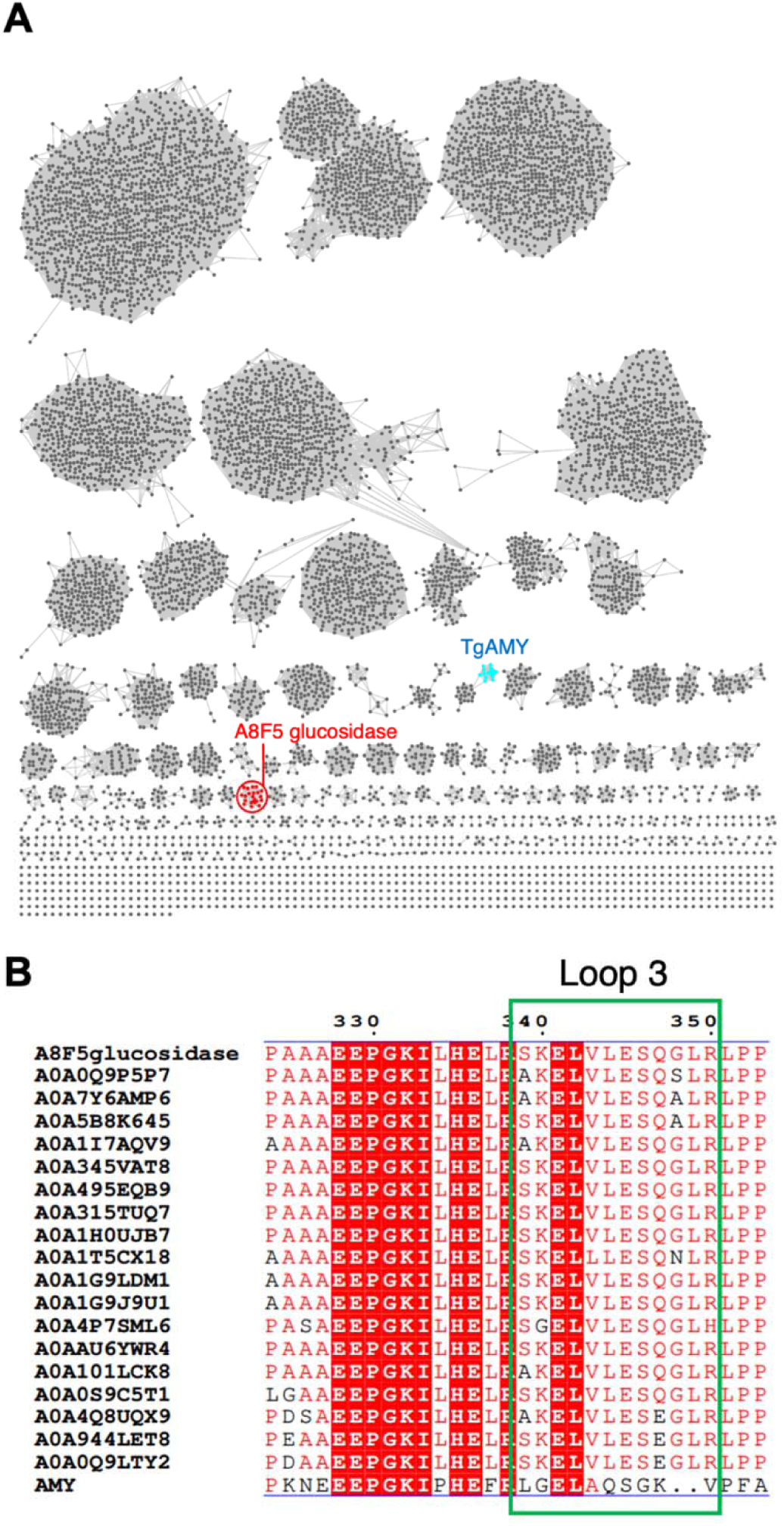
Sequence analysis of A8F5 glucosidase and its homologs in the database. *A*, Sequence similarity network (SSN) of A8F5 glucosidase. Nodes represent 8,177 protein sequences retrieved via a BLAST search using A8F5 glucosidase as the query. Edges connect sequences with an alignment score cutoff of 100. A8F5 glucosidase (red diamond) and AMY (cyan diamond) are partitioned into distinct clusters. *B*, Multiple sequence alignment of the Loop 3 region. All 19 sequences from the A8F5 glucosidase cluster were aligned with the TgAMY sequence using Clustal Omega (52). The flexible Loop 3 region (residues 339–350) is boxed, demonstrating high conservation within the cluster.

## Conclusion

Through the screening of soil bacteria, we discovered A8F5 glucosidase, a kojibiose hydrolase that specifically acts on glucooligosaccharides possessing an α-1,2-linkage and found that it belongs to the GH176 family, which has been scarcely studied to date. The previously known kojibiose hydrolase FjGH65A, also discovered from a soil bacterium genome, possesses a structural architecture in its active site that is distinct from that of A8F5 glucosidase (Fig. 6*A,E*). Consequently, it is inferred that these enzymes acquired the same function through convergent molecular evolution from separate ancestral proteins (clans GH-G and GH-L, respectively).

A8F5 glucosidase possesses four flexible loop regions. In particular, loop 1 is long enough to inhibit crystallization (Fig. 4*A*). Furthermore, the recognition of kojibiose in the active site is surprisingly limited. The interactions with H335 and E336 at subsite +1 are not fully stable, which allows the α-1,2-linked glucoside to occasionally deviate, as observed in the crystal structures (Fig. 5*A*,*B*). This suggests that further tuning through molecular evolution may be required for A8F5 glucosidase to more strictly recognize and hydrolyze kojibiose while maintaining the overall stability and integrity of the protein.

Proteins with amino acid sequences highly similar to that of A8F5 glucosidase were rarely identified even through database searches, and they formed a very small cluster in the SSN analysis (Fig. 7). This indicates that A8F5 glucosidase is an extremely rare entity even within the genomes of highly diverse microbial communities. The rarity of kojibiose hydrolases might correspond to the scarcity of their substrates, kojibiose and kojioligosaccharides, in the environment. Conversely, however, the results of the SSN analysis reveal that multiple clusters of unknown function exist within GH176 alone, suggesting that further genome mining could yield enzymes with novel specificities.

## Experimental procedures

### Assay for kojibiose hydrolyzing activity

Microorganism strains isolated from soil were incubated with reciprocal shaking at 27°C for 3 days in a liquid medium containing 1.5% (w/v) kojibiose, 0.5% (w/v) polypeptone, 0.1% (w/v) yeast extract, 0.1% (w/v) K_2_HPO_4_, 0.06% (w/v) NaH_2_PO_4_·2H_2_O, 0.05% (w/v) MgSO_4_·7H_2_O and 0.3% (w/v) CaCO_3_ (pH 6.8). After cultivation, the culture broth was lysed with 0.1% (v/v) of Triton X-100 and 0.02% (w/v) of lysozyme. Reaction mixtures containing 50 μL of the lysed culture broth and 50 μL of a substrate solution of 4.0% (w/v) kojibiose in 50 mM acetate buffer (pH 6.0) were incubated at 40°C for 48 h. The reaction mixtures were analyzed by TLC. TLC was performed on a Kieselgel 60 plate (Merck, Germany) developed with a solvent system of BuOH:pyridine:water = 6:4:1. Sugar spots were detected by spraying with 20% (v/v) sulfuric acid in MeOH, followed by heating the plates at 120°C for 10 min.

### Strain identification

A single colony of the strain A8F5 was picked, and the DNA was prepared with Mighty prep reagent for DNA (TaKaRa Bio, Kusatsu,, Japan). 16S rRNA was amplified by PCR using the PrimeSTAR Max DNA Polymerase (TaKaRa Bio) with the following primer pair. Forward (8F): 5’-AGAGTTTGATCATGGCTCAG-3’, Reverse (1512R): 5’-TACGGTTACCTTGTTACGACTT-3’. The PCR cycle program was 98°C for 10 s, 50°C for 5 s, and 72°C 10 s for 35 cycles after an initial denaturation at 96°C 2 min. The amplified fragment was sequenced with a DNA sequencer by Eurofins Genomics Co., Ltd. The homologies of the sequences were analyzed according to the NCBI database.

### Enzyme purification from A. humicola A8F5

*A. humicola* A8F5 was cultured at 27°C on a rotary shaker for 3 days in a liquid medium composed of 1.5% (w/v) Pinedex #4, 1.5% (w/v) polypeptone, 0.1% (w/v) yeast extract, 0.1% (w/v) K_2_HPO_4_, 0.06% (w/v) NaH_2_PO_4_·2H_2_O, 0.05% (w/v) MgSO_4_·7H_2_O (pH 6.8). After removal of the supernatant of the culture broth by centrifugation at 10,000 × g for 20 min, the cells were lysed with 0.06% of lysozyme in 10 mM sodium phosphate buffer (pH 7.0). The supernatant of the lysed solution obtained by centrifugation from *A. humicola* A8F5 was collected and precipitated by 80% saturation of ammonium sulfate and stored overnight. The precipitate was collected by centrifugation at 12,000 rpm for 30 min and dissolved in 10 mM sodium phosphate buffer (pH 7.0). After dialysis against the same buffer, the enzyme solution was put on a DEAE-Toyopearl 650S (Tosoh, Tokyo) column (Φ1.0×9.0 cm) equilibrated with the same buffer. Proteins adsorbed on the column were eluted with a linear gradient of 0 to 0.5 M NaCl in the same buffer at a flow rate of 2.0 mL/min. The active fractions were pooled, dialyzed and brought to 1.0 M ammonium sulfate by adding solid ammonium sulfate. The enzyme solution was then loaded onto a Phenyl-Toyopearl 650M (Tosoh) column (Φ1.6×10.0 cm) equilibrated with 10 mM sodium phosphate buffer (pH 7.0) containing 1.0 M ammonium sulfate. The adsorbed proteins were eluted with a linear gradient of 1.0 to 0.5 M ammonium sulfate in the same buffer at a flow rate of 0.5 mL/min. The active fractions were pooled and dialyzed against the same buffer. The enzyme solution was then loaded onto a Resource Q (GE Healthcare) column (Φ0.64×3 cm) equilibrated with 10 mM sodium phosphate buffer (pH 7.0). Proteins adsorbed on the column were eluted with a linear gradient of 0 to 0.5 M NaCl in the same buffer at a flow rate of 0.5 mL/min. The active fractions were pooled, dialyzed and brought to 0.8 M ammonium sulfate by adding solid ammonium sulfate. The enzyme solution was then loaded onto a Butyl-Toyopearl 650M (Tosoh) column (Φ1.6×10.0 cm) equilibrated with 10 mM sodium phosphate buffer (pH 7.0) containing 0.8 M ammonium sulfate. The adsorbed proteins were eluted with a linear gradient of 1.0 to 0.4 M ammonium sulfate in the same buffer at a flow rate of 0.5 mL/min. The active fractions were pooled and dialyzed against the same buffer. The enzyme solution was then loaded onto a Hydroxyapatite (Wako) column (Φ0.5×5.0 cm) equilibrated with 10 mM sodium phosphate buffer (pH 7.0). Non-absorbed proteins were collected as the active fractions. The enzyme was concentrated by using an ultrafiltration centrifugal membrane unit (Amicon Ultra 10 kDa molecular weight cutoff; Merck, Germany), and loaded onto a Superdex 200pg (GE Healthcare, USA) column (Φ2.6×60 cm) with the 10 mM sodium phosphate buffer (pH 7.0) containing 0.4 M NaCl at a flow rate of 2.0 mL/min. The active fractions were pooled and dialyzed against the 10 mM sodium phosphate buffer (pH 7.0). The resulting enzyme solution was used for analyzing the N-terminal amino acid sequence of the enzyme.

### Whole genome analysis of A. humicola A8F5 and gene cloning

The genomic DNA of *A. humicola* A8F5 was extracted using a NucleoBond HMW DNA kit (MACHEREY-NAGEL, Germany) according to the manufacturer’s instructions. The genomic DNA was subjected to high-throughput sequencing using Oxford Nanopore MinION platforms. The N-terminal amino acid sequence of the enzyme was searched from whole-genome sequence, and only one region was matched. To obtain the target gene, PCR was performed using PrimeSTAR Max DNA Polymerase (TaKaRa Bio). The genomic DNA of *A. humicola* A8F5 obtained as above was used as the template for the PCR. The sequence of the forward primer was 5’-TTAAGAAGGAGATATACATATGACTGTTCAGCCAGGCCTGC-3’ and the sequence of the reverse primer was 5’-ATGCTAGCCATACCATGTTAATGATGATGATGATGATGGAGCCCGACGCCCATCGCC-3’. The reverse primer was designed to express C-terminally His_6_-tagged protein. The temperature program for each cycle was 98°C for 10 s, 66°C for 5 s, and 72°C for 10 s. After heat treatment at 95°C for 2 min for DNA denaturation, 35 cycles were run. The products were purified by ethanol precipitation. To gene expression vector for *E. coli* was constructed by inserting the target gene into pRSET A vector (Thermo Fisher, USA) using In-Fusion HD Cloning Kit (TaKaRa Bio) according to the manufacturer’s instructions. The constructed expression plasmid was used to transform *E. coli*. The transformed *E. coli* BL21 (DE3) pLysS (TaKaRa Bio) was cultured in Terrific broth (Thermo Fisher, USA) medium containing antibiotics (50 mg/L ampicillin and 10 mg/L chloramphenicol) at 27°C for 16 h. The cells were harvested by centrifugation at 10,000 rpm for 10 min at 4°C and suspended in a solution of cOmplete Protease Inhibitor Cocktail Tablets (Merck, Germany). To obtain cell-free extracts, the suspended solution was sonicated and centrifuged at 18,000 rpm for 10 min at 4°C. The supernatant was filtered using a 0.45-μm filter and equivalently mixed with 20 mM sodium phosphate buffer (pH 7.4) containing 500 mM NaCl and 40 mM imidazole. The His-tagged protein was purified by nickel affinity column chromatography using Ni Sepharose 6 Fast Flow (GE Healthcare, USA) with two elution steps of 45.5 mM and 150 mM imidazole in 20 mM sodium phosphate buffer (pH 7.4). After buffer exchange by using an ultrafiltration centrifugal membrane unit (Amicon Ultra 50 kDa molecular weight cutoff; Merck, Germany), the concentrated purified His-tagged recombinant protein was used for the enzyme assay.

### Enzyme assay

Hydrolytic activity against kojibiose was assayed as follows. The enzyme solution (0.1 mL) was added to a substrate solution (1.0 mL) containing 0.1% kojibiose in 25 mM sodium phosphate buffer (pH 6.5). After incubation at 40°C for 30 min, the reaction was stopped by boiling the mixture for 10 min, and released glucose was quantitated with the Glucose Assay Kit, Glucose C -test Wako (Wako Pure Chemicals, Japan). One unit (U) of the enzyme activity was defined as the amount of enzyme producing 2 μmol of glucose per min under the stated conditions.

### Effect of temperature, pH and metal ions on activity

The effect of pH on the enzyme activity was examined using Britton-Robinson buffer at 40°C, and the effect of temperature was assayed at 25–75°C using 25 mM sodium phosphate buffer pH 6.5. pH stability was determined by preincubating the enzyme solution at various pH values using 50 mM Britton-Robinson buffer at 4°C for 24 h and measuring the residual activity under the optimum conditions (25 mM sodium phosphate buffer pH 6.5 at 40°C). Thermal stability was determined by preincubating the enzyme at temperatures ranging from 25 to 70°C for 1 h in 25 mM sodium phosphate buffer pH 6.5 and the residual activity was measured under the optimal condition (pH 6.5, 40°C). To determine the effects of various metal ions on the enzyme activity, reactions were performed in the presence of each agent (1 mM) under the optimal condition (pH 6.5, 40°C).

### Substrate specificity of the enzyme

All substrates were prepared enzymatically. Reaction mixtures of 1% (w/v) substrate and the purified recombinant *A. humicola* A8F5 glucosidase (final concentration 0.169 μg/ml) were incubated at 40°C for 24 h in 25 mM sodium phosphate buffer (pH 6.5). After heat deactivation treatment at 100°C for 10 min to stop the reaction, the reaction mixtures were analyzed by TLC as mentioned above or HPLC. Because nigerose is degraded by heat treatment under neutral pH conditions, the reaction was stopped by adding 1.0 M HCl and neutralized by 1.0 M NaOH when nigerose was used as a substrate. The amounts of glucose released by hydrolysis were determined by HPLC. Samples were first treated by filtration using a Millex-LG (0.45 μm, Merck Millipore, Germany) and by deionization using a micro acilyzer G0 (Asahi Chemical Co., Tokyo, Japan). An MCI GEL CK04SS column (10 mm i.d. × 200 mm × 2, Mitsubishi Chem. Co., Tokyo, Japan) was used in the HPLC analysis at a flow rate of 0.4 mL/min at 80°C using water as a solvent. Kinetic parameters of the enzyme activity for kojibiose and nigerose were measured using kojibiose (0.21–13.4 mM) or nigerose (0.17–133 mM) under the optimal conditions (pH 6.5, 40°C).

### Anomeric configuration analysis of the hydrolysis product

The anomeric configuration of the glucose released from selaginose or kojibiose by A8F5 glucosidase was determined by gas-liquid chromatography. The reaction products of 4-*O*-α-glucosyl trehalose by anomer-retaining α-glucosidase (transglucosidase L, Amano Enzyme Inc., Kagamihara, Japan) and anomer-inverting glucoamylase (glucoamylase XL-4, Nagase Viita Corp., Okayama, Japan) were also analyzed as references. The reaction mixture, consisting of 500 μL of 6.0 mM substrate solution, 20 μL of 50 mM sodium phosphate buffer (pH 6.5), and 50 μL of 50 U/mL enzyme solution, was incubated at room temperature. Aliquots of 50 μL of the reaction mixture were pipetted out at appropriate times and transferred into test tubes for the determination of glucose and the anomeric form. Each aliquot for the analysis of anomeric forms was immediately frozen by immersing the test tube in a cold ethanol bath and then lyophilized for 1 h. The freeze-dried sample was converted to the trimethylsilyl compound by incubation with 100 μL of trimethylsilylation reagent (Tokyo Chemical Industry, Japan) at 60°C for 30 min and subjected to gas-liquid chromatography. Gas-liquid chromatography was performed on a Shimadzu Model 2010 Plus gas-liquid chromatograph equipped with a hydrogen flame-ionization detector. A ZB-50 column (30 m×0.25 mm, Phenomenex) was employed. Helium was used as the carrier gas at a flow rate of 35 cm/sec.

### Protein purification and crystallography

For preparation of a protein sample for crystallization with an N-terminal His_6_-tag, the expression vector was changed to pET28a (Novagen, Maidon, WI, USA) by using In-Fusion HD Cloning Kit (TaKaRa Bio) with the following primer pair. Forward: 5’-CATCATCATCACATGACTGTTCAG-3’, Reverse: 5’-TTCGGGCTTTGTTAGAGCCCG-3’. The overexpression plasmid for the D427N mutant was constructed by site-directed mutagenesis using KOD One PCR Master Mix (TOYOBO Co., Ltd., Osaka, Japan) and methylation-sensitive restriction enzyme DpnI (TaKaRa Bio). The following primers were used: 5’-CCAGGGCTGGAAG<u>AAT</u>TCCCGGGATGC-3’ (forward) and 5’-GCATCCCGGGA<u>ATT</u>CTTCCAGCCCTGG-3’ (reverse), where the mutated codon is underlined. *E. coli* BL21-CodonPlus (DE3)-RIL (Agilent Technologies, Santa Clara, CA, USA) harboring the overexpression plasmid was cultured in lysogeny broth medium containing 50 mg/L kanamycin and 17 mg/L chloramphenicol at 37 °C until the OD600 reached approximately 0.5. Protein expression was induced with 0.1 mM isopropyl-β-D-thiogalactopyranoside at 25°C for 20 h. The cells were harvested by centrifugation and suspended in 50 mM Tris-HCl (pH 7.5) and 300 mM NaCl (lysis buffer). Cell extracts were obtained by sonication, followed by centrifugation to remove cell debris. The supernatant was filtered using a Millex-HP Syringe Filger Unit with 0.45 μm polyethersulfone membrane (Millipore, Billerica, MA, USA). The crude enzyme was applied to a Ni-NTA agarose column (Qiagen, Hilden, Germany) equilibrated with the lysis buffer. The column was washed with the lysis buffer containing 20 mM imidazole, and the His-tagged protein was eluted with the lysis buffer containing 500 mM imidazole. After elution, proteins were concentrated and buffer-exchanged using Amicon Ultra-15 Centrifugal Filter Devices (MWCO 10 kDa, Millipore). The enzyme was further purified by gel filtration column chromatography using a Superose 6 10/300 GL column (Cytiva, Marlborough, MA, USA) equilibrated with 20 mM Tris-HCl (pH 7.5) and 300 mM NaCl. The purified sample was desalted and concentrated using an Amicon Ultra-15 Centrifugal Filter Devices (MWCO 10 kDa, Millipore) into 20 mM Tris-HCl (pH 7.5) and 3 mM NaCl.

Crystallization was performed using the sitting-drop vapor-diffusion method by mixing equal volumes (0.5 μL each) of a protein solution (15 mg/mL WT A8F5 glucosidase or 10 mg/mL Δloop1) and a reservoir solution consisting of 0.1 M sodium acetate (pH 5.1) and 34% (v/v) 1,2-propanediol. For the glucose-complex structure, co-crystallization was performed by incubating the protein solution with 10 mM glucose prior to setting up the drops. For other substrate-complex structures, apo-crystals were soaked for 1–24 h in a modified reservoir solution containing 10–20 mM of each ligand (kojibiose, kojitriose, or selaginose). The crystals were cryoprotected by 20% (v/v) PEG 200, harvested using nylon loops, and flash-cooled in liquid nitrogen. X-ray diffraction data were collected at beamlines AR-NE3A, BL-1A and BL-5A of the Photon Factory (Tsukuba, Ibaraki, Japan) and BL45XU of SPring-8 (Sayo, Hyogo, Japan). The data were automatically processed, integrated, and scaled using XDS (41). Subsequent structural analysis was performed using the CCP4 suite (42). The structure of A8F5 glucosidase was determined by molecular replacement with PHASER (43), using the structure predicted by ColabFold (44) as a search template. Iterative cycles of manual model building and refinement were performed using COOT (45) and REFMAC5 (46), respectively. The final model was validated using MolProbity (47). All structural figures were prepared using UCSF Chimera (48) and PyMOL (Shrödinger LLC). Polder maps were created using the PHENIX software (49).

### Sequence similarity network (SSN)

An SSN was constructed using the EFI Enzyme Similarity Tool (EFI-EST) (50). The amino acid sequence of A8F5 glucosidase was used as a query to retrieve related sequences from the UniProtKB database, with an initial E-value threshold of 10^-1^. An alignment score threshold of 100 was applied to the resulting network, such that edges were drawn only between protein sequences sharing a similarity score of 100 or greater. The network was visualized and analyzed using Cytoscape (51), where each node represents a protein sequence and each edge represents a pairwise similarity above the specified threshold.

## Data availability

The nucleotide sequence of the A8F5 glucosidase gene has been deposited in the DDBJ/ENA/GenBank databases under accession number LC922693. The atomic coordinates and structure factors (PDB codes: 24AI, 24AZ, 24IL, 24IR, and 24IS) have been submitted to the Protein Data Bank (https://wwpdb.org/).

## Acknowledgments

We thank Ms. Mayuko Matsumoto for her efforts in the preparation of the expression vector and initial crystallization screening, and Mr. Noriaki Kitagawa, Mr. Hiroki Asakuma, Dr. Masahiro Sota, and Mr. Takumi Masaki for their assistance in promoting collaborative research. We also thank Drs. Takatoshi Arakawa, Chihaya Yamada, Hajime Aga, Shimpei Ushio, and Koryu Yamamoto for their invaluable technical assistance and insightful discussions. We acknowledge the staff at SPring-8 and the Photon Factory for their support with X-ray data collection.

## Author contributions

T. M., H. W., and S. F. conceptualization; T. K., A. M., and S. F. data curation; R. Y., T. S., and S. F. formal analysis; T. M., A. M., H. W., and S. F. funding acquisition; R. Y., T. S., T. T., and K. H., investigation; T. K., and A. M. methodology; T. M., T. K., and A. M. project administration; H. W. and S. F. resources; H. W. and S. F. supervision; S. F. validation; R. Y. and T. S. visualization; R. Y., T. S., and S. F. writing – original draft; R. Y., T. S., T. T., K. H., T. M., T. K., A. M., H. W., and S. F. writing – review & editing.

## Funding and additional information

This work was partially supported by JSPS-KAKENHI (24H02269, 23H00322, and 23K21168 to SF), the Research Support Project for Life Science and Drug Discovery (Basis for Supporting Innovative Drug Discovery and Life Science Research (BINDS)) from AMED under Grant Numbers JP22ama121001, JP23ama121001, and JP24ama121001.

## Abbreviations

GH: glycoside hydrolase
TLC: thin-layer chromatography
ORF: open reading frame
TgAMY: *Thermococcus gammatolerans* GH176 amylo-α-1,6-glucosidase
HPLC: high-performance liquid chromatography
tNCS: translational non-crystallographic symmetry
pLDDT: predicted local distance test
RMSD: root mean square deviations
FjGH65A: *Flavobacterium johnsoniae* GH65 kojibiose hydrolase
SSN: sequence similarity network.

---

**Table S1.** Effects of metal salts on the A8F5 glucosidase activity.

| **Metal salts** | **Relative activity (%)** | **Metal salts** | **Relative activity (%)** |
| --- | --- | --- | --- |
| None | 100.0 | MnCl<sub>2</sub> | 100.1 |
| EDTA | 95.5 | CoCl<sub>2</sub> | 96.1 |
| RbCl | 99.1 | NiCl<sub>2</sub> | 86.1 |
| KCl | 97.9 | PbCl<sub>2</sub> | 101.0 |
| NaCl | 100.9 | SnCl<sub>2</sub> | 97.5 |
| NH<sub>4</sub>Cl | 100.6 | SrCl<sub>2</sub> | 100.6 |
| LiCl | 101.0 | CaCl<sub>2</sub> | 101.5 |
| MgCl<sub>2</sub> | 100.6 | FeCl<sub>2</sub> | 96.2 |
| BaCl<sub>2</sub> | 102.1 | ZnCl<sub>2</sub> | 88.2 |
| HgCl<sub>2</sub> | 1.2 | AlCl<sub>3</sub> | 97.2 |
| CuCl<sub>2</sub> | 3.4 | FeCl<sub>3</sub> | 95.1 |

Enzyme activity was measured in the presence of metal salts or a reagent (1 mM) under the optimal condition (pH 6.5, 40°C). EDTA: ethylenediaminetetraacetic acid.

**Table S2.**
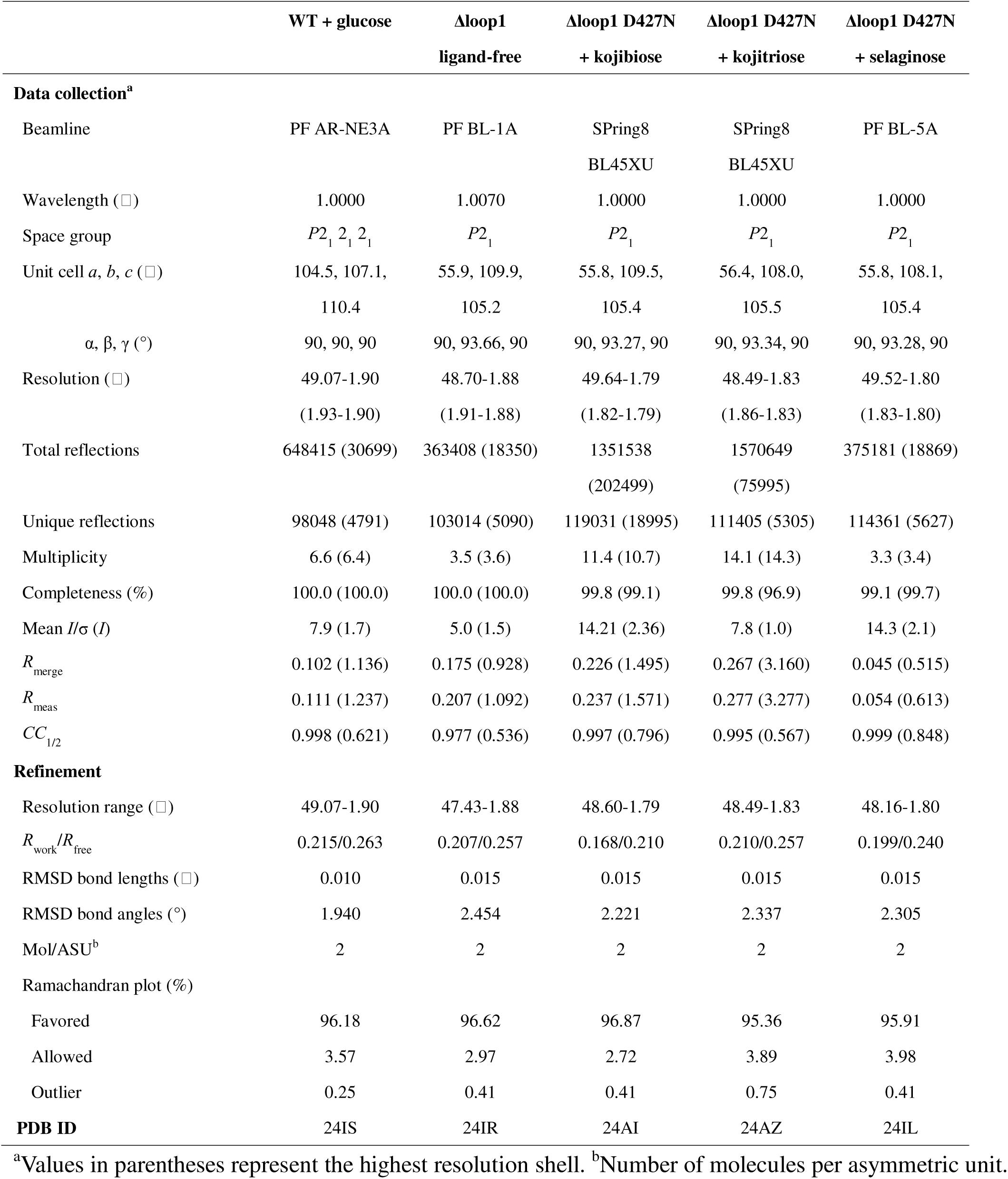
Crystallographic data collection and refinement statistics of A8F5 glucosidase.

**Table S3.**
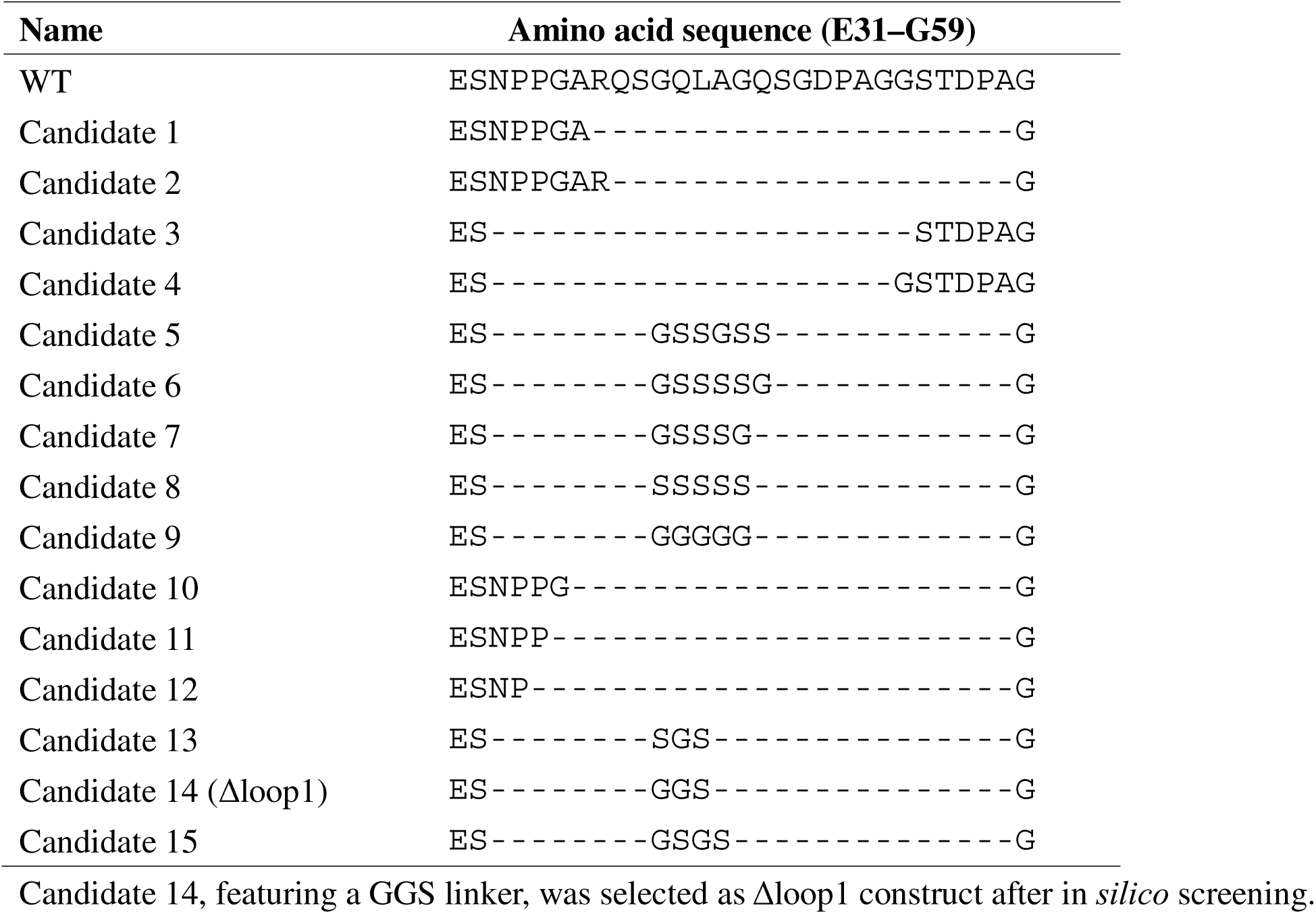
Designed loop deletants.

**Figure S1.**
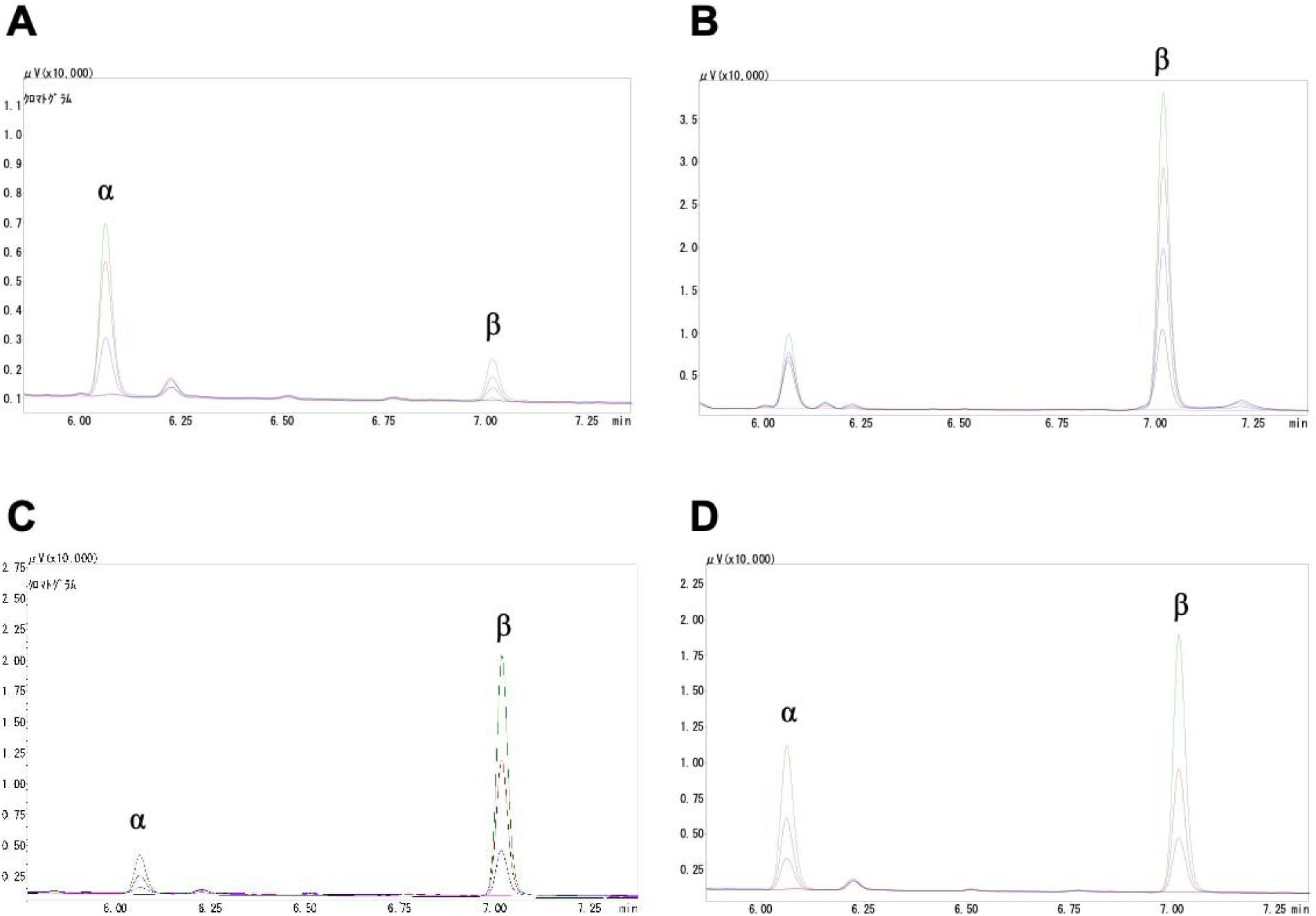
Analysis of hydrolysis products. The increasing time course of each glucose anomer after the reaction was analyzed by gas-liquid chromatography. *A*, Reaction of 4-*O*-α-glucosyl trehalose with an anomer-retaining α-glucosidase. *B*, Reaction of 4-*O*-α-glucosyl trehalose with an anomer-inverting glucoamylase. *C*, Reaction of selaginose with A8F5 glucosidase. *D*, Reaction of kojibiose with A8F5 glucosidase. Chromatograms around the glucose peaks for the substrate without enzyme (pink), enzyme without substrate (black), and reaction products after 0.25 min (blue), 0.75 min (brown), and 1.5 min (green) are shown. Details of the analysis are described in the Experimental Procedures.

**Figure S2.**
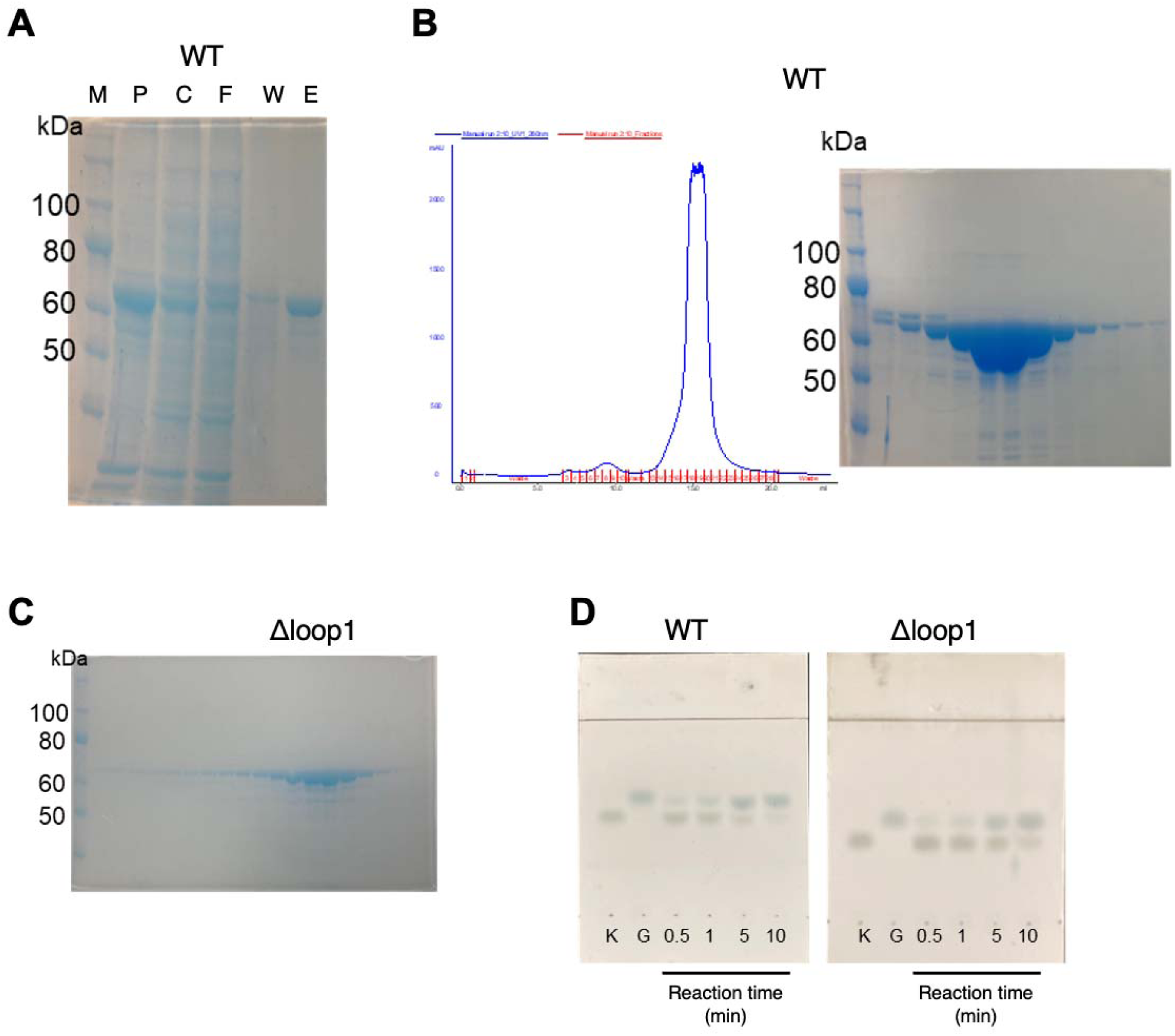
Purification and activity of recombinant proteins of wild-type (WT) A8F5 glucosidase and the Δloop1 construct. *A*, SDS-PAGE of each fraction before and after nickel affinity chromatography during WT protein purification. Symbols: M, molecular size marker; P, pellet; C, crude supernatant; F, flow-through; W, wash; E, eluate. *B*, Gel filtration chromatography for WT protein purification. The chromatogram (left) and SDS-PAGE of the collected eluate fractions (right) are shown. *C*, SDS-PAGE of the collected eluate fractions from gel filtration chromatography of Δloop1. *D*, TLC analysis on the activity of WT A8F5 glucosidase (left) and Δloop1 (right). Symbols: K, 1 mM kojibiose; G, 1 mM glucose; 0.5–10, reaction time (min).

## References

1. Cummings, R. D. (2024) A periodic table of monosaccharides. Glycobiology. 34, cwad088

2. Lombard, V., Henrissat, B., and Garron, M. L. (2025) CAZac: an activity descriptor for carbohydrate-active enzymes. Nucleic Acids Res. 53, D625–D633

3. Overkleeft, H. S., Davies, G. J., and Williams, S. J. (2026) Catalyzing carbohydrate cleavage: Glycosidases and their mechanisms. Chem. Rev. 126, 3287–3323

4. Zhang, S., Ni, D., Zhu, Y., Xu, W., Zhang, W., and Mu, W. (2024) A comprehensive review on the properties, production, and applications of functional glucobioses. Crit. Rev. Food Sci. Nutr. 64, 13149–13162

5. Elango, B., Shirley, C. P., Okram, G. S., Ramesh, T., Seralathan, K. K., and Mathanmohun, M. (2023) Structural diversity, functional versatility and applications in industrial, environmental and biomedical sciences of polysaccharides and its derivatives – A review. Int. J. Biol. Macromol. 250, 126193

6. Ruiz-Herrera, J., and Ortiz-Castellanos, L. (2019) Cell wall glucans of fungi. A review. The Cell Surface. 5, 100022

7. Matsuda, K. (1957) Kojibiose (2-O-α-D-glucopyranosyl-D-glucose): Isolation and structure: Chemical synthesis. Nature 1957 180:4593. 180, 985–985

8. Zhou, Z., Zhang, H., Lou, T., Wang, S., Fan, Y., Niu, S., and Ge, L. (2026) Research progress in kojibiose: structural insights, enzymatic synthesis, and applications in food and pharmaceutical industries. Crit. Rev. Food Sci. Nutr. 10.1080/10408398.2025.2610756

9. Ogawa, S., Ashiura, M., and Uchida, C. (1998) Synthesis of α-glucosidase inhibitors: kojibiose-type pseudodisaccharides and a related pseudotrisaccharide. Carbohydr. Res. 307, 83–95

10. McGrath, D., Lee, E. E., and O’Colla, P. S. (1969) The chemical synthesis of polysaccharides : Part II. (1→2)-, (1→3)-, and (1→4)-linked glucans. Carbohydr. Res. 11, 461–465

11. Chaen, H., Nishimoto, T., Nakada, T., Fukuda, S., Kurimoto, M., and Tsujisaka, Y. (2001) Enzymatic synthesis of novel oligosaccharides from L-sorbose, maltose, and sucrose using kojibiose phosphorylase. J. Biosci. Bioeng. 92, 173–176

12. Verhaeghe, T., De Winter, K., Berland, M., De Vreese, R., D’Hooghe, M., Offmann, B., and Desmet, T. (2016) Converting bulk sugars into prebiotics: semi-rational design of a transglucosylase with controlled selectivity. Chemical Communications. 52, 3687–3689

13. Chaen, H., Yamamoto, T., Nishimoto, T., Nakada, T., Fukuda, S., Sugimoto, T., Kurimoto, M., and Tsujisaka, Y. (1999) Purification and characterization of a novel phosphorylase, kojibiose phosphorylase, from *Thermoanaerobium brockii*. J. Appl. Glycosci. (1999). 46, 423–429

14. Barker, M. K., and Rose, D. R. (2013) Specificity of processing α-glucosidase I is guided by the substrate conformation: Crystallographic and in silico studies. J. Biol. Chem. 288, 13563–13574

15. Hamazaki, H., and Hotta, K. (1980) Enzymatic hydrolysis of disaccharide unit of collagen. Eur. J. Biochem. 111, 587–591

16. Kobayashi, M., Mitsuishi, Y., and Matsuda, K. (1978) Pronounced hydrolysis of highly branched dextrans with a new type of dextranase. Biochem. Biophys. Res. Commun. 80, 306–312

17. Miyazaki, T., Tanaka, H., Nakamura, S., Dohra, H., and Funane, K. (2023) Identification and characterization of dextran α-1,2-debranching enzyme from *Microbacterium dextranolyticum*. J. Appl. Glycosci. 70, 15–24

18. Nakamura, S., Nihira, T., Kurata, R., Nakai, H., Funane, K., Park, E. Y., and Miyazaki, T. (2021) Structure of a bacterial α-1,2-glucosidase defines mechanisms of hydrolysis and substrate specificity in GH65 family hydrolases. J. Biol. Chem. 297, 101366

19. De Beul, E., Jongbloet, A., Franceus, J., and Desmet, T. (2021) Discovery of a kojibiose hydrolase by analysis of specificity-determining correlated positions in glycoside hydrolase family 65. Molecules. 26, 6321

20. Drula, E., Garron, M. L., Dogan, S., Lombard, V., Henrissat, B., and Terrapon, N. (2022) The carbohydrate-active enzyme database: functions and literature. Nucleic Acids Res. 50, D571–D577

21. Wang, Y., Tian, Y., Ban, X., Li, C., Hong, Y., Cheng, L., Gu, Z., and Li, Z. (2022) Substrate selectivity of a novel amylo-α-1,6-glucosidase from *Thermococcus gammatolerans* STB12. Foods 2022, Vol. 11, Page 1442. 11, 1442

22. Chalita, M., Kim, Y. O., Park, S., Oh, H. S., Cho, J. H., Moon, J., Baek, N., Moon, C., Lee, K., Yang, J., Nam, G. G., Jung, Y., Na, S. I., Bailey, M. J., and Chun, J. (2024) EzBioCloud: a genome-driven database and platform for microbiome identification and discovery. Int. J. Syst. Evol. Microbiol. 74, 006421

23. Kageyama, A., Morisaki, K., Omura, S., and Takahashi, Y. (2008) *Arthrobacter oryzae* sp. nov. and *Arthrobacter humicola* sp. nov. Int. J. Syst. Evol. Microbiol. 58, 53–56

24. Holm, L. (2020) DALI and the persistence of protein shape. Protein Sci. 29, 128–140

25. Tarkowski, L. P., Tsirkone, V. G., Osipov, E. M., Beelen, S., Lammens, W., Vergauwen, R., Van Den Ende, W., and Strelkov, S. V. (2020) Crystal structure of *Arabidopsis thaliana* neutral invertase 2. Acta Crystallogr. Sect. F Struct. Biol. Cryst. Commun. 76, 152–157

26. Xie, J., Cai, K., Hu, H. X., Jiang, Y. L., Yang, F., Hu, P. F., Cao, D. D., Li, W. F., Chen, Y., and Zhou, C. Z. (2016) Structural analysis of the catalytic mechanism and substrate specificity of *Anabaena* alkaline invertase InvA reveals a novel glucosidase. J. Biol. Chem. 291, 25667–25677

27. Miyazaki, T., Ichikawa, M., Yokoi, G., Kitaoka, M., Mori, H., Kitano, Y., Nishikawa, A., and Tonozuka, T. (2013) Structure of a bacterial glycoside hydrolase family 63 enzyme in complex with its glycosynthase product, and insights into the substrate specificity. FEBS J. 280, 4560–4571

28. Gibson, R. P., Gloster, T. M., Roberts, S., Warren, R. A. J., Storch De Gracia, I., García, Á., Chiara, J. L., and Davies, G. J. (2007) Molecular basis for trehalase inhibition revealed by the structure of trehalase in complex with potent inhibitors. Angew. Chem. Int. Ed. 46, 4115–4119

29. Mhlongo, N. N., Skelton, A. A., Kruger, G., Soliman, M. E. S., and Williams, I. H. (2014) A critical survey of average distances between catalytic carboxyl groups in glycoside hydrolases. Proteins: Structure, Function and Bioinformatics. 82, 1747–1755

30. Lovelace, J. J., and Borgstahl, G. E. O. (2020) Characterizing pathological imperfections in macromolecular crystals: lattice disorders and modulations. Crystallogr. Rev. 26, 3–50

31. Abramson, J., Adler, J., Dunger, J., Evans, R., Green, T., Pritzel, A., Ronneberger, O., Willmore, L., Ballard, A. J., Bambrick, J., Bodenstein, S. W., Evans, D. A., Hung, C. C., O’Neill, M., Reiman, D., Tunyasuvunakool, K., Wu, Z., Žemgulytė, A., Arvaniti, E., Beattie, C., Bertolli, O., Bridgland, A., Cherepanov, A., Congreve, M., Cowen-Rivers, A. I., Cowie, A., Figurnov, M., Fuchs, F. B., Gladman, H., Jain, R., Khan, Y. A., Low, C. M. R., Perlin, K., Potapenko, A., Savy, P., Singh, S., Stecula, A., Thillaisundaram, A., Tong, C., Yakneen, S., Zhong, E. D., Zielinski, M., Žídek, A., Bapst, V., Kohli, P., Jaderberg, M., Hassabis, D., and Jumper, J. M. (2024) Accurate structure prediction of biomolecular interactions with AlphaFold 3. Nature. 630, 493–500

32. Aleshin, A. E., Stoffer, B., Firsov, L. M., Svensson, B., and Honzatko, R. B. (1996) Crystallographic complexes of glucoamylase with maltooligosaccharide analogs: Relationship of stereochemical distortions at the nonreducing end to the catalytic mechanism. Biochemistry. 35, 8319–8328

33. Hidaka, M., Kitaoka, M., Hayashi, K., Wakagi, T., Shoun, H., and Fushinobu, S. (2006) Structural dissection of the reaction mechanism of cellobiose phosphorylase. Biochem. J. 398, 37–43

34. Okada, S., Yamamoto, T., Watanabe, H., Nishimoto, T., Chaen, H., Fukuda, S., Wakagi, T., and Fushinobu, S. (2014) Structural and mutational analysis of substrate recognition in kojibiose phosphorylase. FEBS J. 281, 778–786

35. Hasegawa, K., Kubota, M., and Matsuura, Y. (1999) Roles of catalytic residues in α-amylases as evidenced by the structures of the product-complexed mutants of a maltotetraose-forming amylase. Protein Eng. 12, 819–824

36. Matsuura, Y. (2002) A possible mechanism of catalysis involving three essential residues in the enzymes of α-amylase family. Biologia (Bratislava). 57, 21–27

37. Castro, N., Durrieu, V., Raynaud, C., and Rouilly, A. (2016) Influence of DE-value on the physicochemical properties of maltodextrin for melt extrusion processes. Carbohydr. Polym. 144, 464–473

38. Woo, E. J., Lee, S., Cha, H., Park, J. T., Yoon, S. M., Song, H. N., and Park, K. H. (2008) Structural insight into the bifunctional mechanism of the glycogen-debranching enzyme TreX from the Archaeon Sulfolobus solfataricus. Journal of Biological Chemistry. 283, 28641–28648

39. Song, H. N., Jung, T. Y., Park, J. T., Park, B. C., Myung, P. K., Boos, W., Woo, E. J., and Park, K. H. (2010) Structural rationale for the short branched substrate specificity of the glycogen debranching enzyme GlgX. Proteins: Structure, Function, and Bioinformatics. 78, 1847–1855

40. Sim, L., Beeren, S. R., Findinier, J., Dauville, D., Ball, S. G., Henriksen, A., and Palcic, M. M. (2014) Crystal structure of the *Chlamydomonas* starch debranching enzyme isoamylase ISA1 reveals insights into the mechanism of branch trimming and complex assembly. J. Biol. Chem. 289, 22991–23003

41. Kabsch, W. (2010) XDS. Acta Crystallogr. Sect. D-Struct. Biol. 66, 125–132

42. Potterton, L., Agirre, J., Ballard, C., Cowtan, K., Dodson, E., Evans, P. R., Jenkins, H. T., Keegan, R., Krissinel, E., Stevenson, K., Lebedev, A., McNicholas, S. J., Nicholls, R. A., Noble, M., Pannu, N. S., Roth, C., Sheldrick, G., Skubak, P., Turkenburg, J., Uski, V., Von Delft, F., Waterman, D., Wilson, K., Winn, M., and Wojdyr, M. (2018) CCP4i2: The new graphical user interface to the CCP4 program suite. Acta Crystallogr. D Struct. Biol. 74, 68–84

43. McCoy, A. J., Grosse-Kunstleve, R. W., Adams, P. D., Winn, M. D., Storoni, L. C., and Read, R. J. (2007) Phaser crystallographic software. J. Appl. Crystallogr. 40, 658–674

44. Mirdita, M., Schütze, K., Moriwaki, Y., Heo, L., Ovchinnikov, S., and Steinegger, M. (2022) ColabFold: making protein folding accessible to all. Nat. Methods. 19, 679–682

45. Emsley, P., Lohkamp, B., Scott, W. G., and Cowtan, K. (2010) Features and development of Coot. Acta Crystallogr. Sect. D-Struct. Biol. 66, 486–501

46. Murshudov, G. N., Skubák, P., Lebedev, A. A., Pannu, N. S., Steiner, R. A., Nicholls, R. A., Winn, M. D., Long, F., and Vagin, A. A. (2011) REFMAC5 for the refinement of macromolecular crystal structures. Acta Crystallogr. D Biol. Crystallogr. 67, 355–367

47. Williams, C. J., Headd, J. J., Moriarty, N. W., Prisant, M. G., Videau, L. L., Deis, L. N., Verma, V., Keedy, D. A., Hintze, B. J., Chen, V. B., Jain, S., Lewis, S. M., Arendall, W. B., Snoeyink, J., Adams, P. D., Lovell, S. C., Richardson, J. S., and Richardson, D. C. (2018) MolProbity: More and better reference data for improved all-atom structure validation. Protein Science. 27, 293–315

48. Pettersen, E. F. F., Goddard, T. D. D., Huang, C. C. C., Couch, G. S. S., Greenblatt, D. M. M., Meng, E. C. C., and Ferrin, T. E. E. (2004) UCSF Chimera - A visualization system for exploratory research and analysis. J. Comput. Chem. 25, 1605–1612

49. Liebschner, D., Afonine, P. V., Moriarty, N. W., Poon, B. K., Sobolev, O. V., Terwilliger, T. C., and Adams, P. D. (2017) Polder maps: Improving OMIT maps by excluding bulk solvent. Acta Crystallogr. D Struct. Biol. 73, 148–157

50. Zallot, R., Oberg, N., and Gerlt, J. A. (2019) The EFI web resource for genomic enzymology tools: Leveraging protein, genome, and metagenome databases to discover novel enzymes and metabolic pathways. Biochemistry. 58, 4169–4182

51. Shannon, P., Markiel, A., Ozier, O., Baliga, N. S., Wang, J. T., Ramage, D., Amin, N., Schwikowski, B., and Ideker, T. (2003) Cytoscape: A software environment for integrated models of biomolecular interaction networks. Genome Res. 13, 2498–2504

52. Sievers, F., Wilm, A., Dineen, D., Gibson, T. J., Karplus, K., Li, W., Lopez, R., McWilliam, H., Remmert, M., Söding, J., Thompson, J. D., and Higgins, D. G. (2011) Fast, scalable generation of high-quality protein multiple sequence alignments using Clustal Omega. Mol. Syst. Biol. 7, 539

